# MicroRNA775 Promotes Intrinsic Leaf Size and Reduces Cell Wall Pectin Level via a Target Galactosyltransferase in *Arabidopsis*

**DOI:** 10.1101/2020.09.17.301705

**Authors:** He Zhang, Zhonglong Guo, Yan Zhuang, Yuanzhen Suo, Jianmei Du, Zhaoxu Gao, Jiawei Pan, Li Li, Tianxin Wang, Liang Xiao, Genji Qin, Yuling Jiao, Huaqing Cai, Lei Li

## Abstract

Plants possess unique primary cell walls made of complex polysaccharides that play critical roles in determining intrinsic cell and organ size. How genes responsible for synthesizing and modifying the polysaccharides are regulated by microRNAs (miRNAs) to control plant size remains largely unexplored. Here we identified 23 putative cell wall related miRNAs, termed CW-miRNAs, in *Arabidopsis thaliana* and characterized miR775 as an example. We showed that miR775 post-transcriptionally silences *GALT9*, which encodes an endomembrane-located galactosyltransferase belonging to the glycosyltransferase 31 family. Over-expression of miR775 and deletion of *GALT9* significantly enlarged leaf-related organs, primarily owing to increases in cell size. Monosaccharide quantification, confocal Raman imaging, and immunolabelling combined with atomic force microscopy (AFM) revealed that the *MIR775A*-*GALT9* circuit modulates pectin level and elastic modulus of the cell wall. We further showed that *MIR775A* is directly repressed by the transcription factor ELONGATED HYPOCOTYL 5 (HY5). Genetic analysis confirmed that *HY5* is a negative regulator of leaf size and acts through the *HY5-MIR775A-GALT9* repression cascade to control pectin level. These results demonstrate that miR775-regulated cell wall remodeling is an integral determinant for intrinsic leaf size in *A. thaliana* and highlight the need to study other CW-miRNAs for more insights into cell wall biology.

## Introduction

Precise control of organ size is a fundamental feature of living organisms that results in distinct, species-specific organ sizes and shapes (Bogre et al., 2008; Johnson and Lenhard, 2011; Hong et al., 2018). Genetic analyses in both animals and plants have established that intrinsic organ size is determined by the combinatory effects of cell proliferation and cell expansion (Bogre et al., 2008; Johnson and Lenhard, 2011; Gonzalez et al., 2012; Tumaneng et al., 2012; Hepworth and Lenhard, 2014; Hong et al., 2018). Over the past two decades, an increasingly detailed picture is emerging on cell proliferation control in plants, which involves transcriptional regulators (Mizukami and Fischer, 2000; Powell and Lenhard, 2012; Du et al., 2014), miRNAs (Rodriguez et al., 2010; Schommer et al., 2014; Yang et al., 2018), and the ubiquitin-proteasome pathway (Du et al., 2014). By comparison, our understanding of cell size control in plants is relatively sparse (Ferjani et al., 2007; Hong et al., 2018).

Different from metazoan cells, plant cells are enclosed in the cell walls, which locate between the middle lamella and the plasma membrane. To reach the desired size, plant cells rely on the balance between the inner turgor pressure and the extensibility of the cell walls (Cosgrove, 2005; Palin and Geitmann, 2012; Hong et al., 2018). During growth and development, cell walls need to be loosened in a highly controlled way to allow nondestructive cell expansion, which might increase cell size by several orders of magnitude (Velasquez et al., 2011; Palin and Geitmann, 2012; Hong et al., 2018). Moreover, being sessile organisms, plants are extremely sensitive to the environment and exhibit a number of plastic responses, which allow them to reliably tune size and shape according to the prevailing environmental conditions (Hepworth and Lenhard, 2014; Hong et al., 2018). For example, in response to shading from neighbors, many plants undergo increased stem and petiole elongation in the well-characterized shade avoidance responses. Therefore, the plant cell wall is critical for determining both the intrinsic organ size and how it is shaped by the environment.

Primary plant cell wall is a highly complex and dynamic structure mainly composed of cellulose, hemicelluloses, and pectin (Somerville et al., 2004; Cosgrove, 2005; Somerville, 2006; Palin and Geitmann, 2012). These polysaccharide constituents have different structural and biological roles. Pectin is defined as a group of polysaccharides containing galacturonic acid that acts as gel-forming polymers to cross-link the hemicellulose and cellulose microfibrils (Somerville, 2006; Palin and Geitmann, 2012; Atmodjo et al., 2013). Studies using solid-state nuclear magnetic resonance spectroscopy presented compelling evidence for extensive cellulose-pectin contacts but less cellulose-hemicellulose interactions in the cell walls than previously envisaged (Wang et al., 2015), suggesting that pectin plays an underappreciated role in cell wall remodeling.

Three major classes of pectin polymers have been identified in the cell wall matrix. These include homogalacturonan (HG), which possesses a backbone of 1,4-linked α-D-galacturonosyluronic acid residues, rhamnogalacturonan I (RG-I), which consists of interspersed α-D-galacturonosyl and rhamnosyl residues with galactosyl and arabinosyl side-chains, and the lesser abundant rhamnogalacturonan II (RG-II) (Harholt et al., 2010; Palin and Geitmann, 2012; Atmodjo et al., 2013). Structural data indicate that these pectic constitutes interconnect with each other in the wall via covalent linkages of their backbones (Atmodjo et al., 2013). Recently, nanoimaging studies have showed that HG in pavement cell walls may assemble into discrete nanofilaments rather than an interlinked network (Haas et al., 2020). It was suggested that local and polarized expansion of the HG nanofilaments could lead to cell enlargement without turgor-driven growth (Haas et al., 2020). However, biosynthesis and modifications of the pectin polysaccharides are highly complicated processes and their roles in cell wall remodeling remain to be fully elucidated. Given that the involved enzymes are likely integral membrane proteins in their active forms and the lack of robust *in vivo* assays, functional details of the pectin-related genes in regulating intrinsic organ size remain largely unknown (Qu et al., 2008; Harholt et al., 2010; Palin and Geitmann, 2012; Parsons et al., 2012; Atmodjo et al., 2013; Tan et al., 2013; Qin et al., 2017).

MiRNAs are an endogenous class of sequence-specific, *trans*-acting small regulatory RNAs that modulate gene expression mainly at the post-transcriptional level (Voinnet, 2009; Ma et al., 2010; Yang et al., 2012; Rogers and Chen, 2013). In plants, miRNAs are recognized to regulate an enormous collection of target genes that are implicated in numerous biological processes (Voinnet, 2009; Rogers and Chen, 2013; Rodriguez et al., 2016; Guo et al., 2020). Genetic analysis has uncovered that several miRNAs (e.g. miR319, miR396, and miR408) participate in regulating cell proliferation and organ growth (Palatnik et al., 2003; Ori et al., 2007; Rodriguez et al., 2010; Schommer et al., 2014; Zhang et al., 2014; Rodriguez et al., 2016; Pan et al., 2018; Yang et al., 2018). However, no systematic efforts have been reported to identify and functionally study miRNAs pertinent to the regulation of primary cell wall, even though hundreds of genes are involved in wall biosynthesis and modifications. We reasoned that elucidation of the regulatory roles of cell wall related miRNAs, termed CW-miRNAs, should help expanding our understanding of how cell wall remodeling contributes to intrinsic organ size adjustment in plants.

In the current study, we identified a group of 23 putative CW-miRNAs in *A. thaliana*. We focused on functional characterization of miR775 as an exemplar CW-miRNA and delineated the *HY5-MIR775A-GALT9* repression pathway for modulating cell size and leaf size. Cellular analyses combining monosaccharide quantification, confocal Raman imaging, immunolabelling, and atomic force microscopy (AFM) revealed that this pathway regulates pectin level and elastic modulus of the cell wall. Collectively, these results demonstrated the importance of miRNA-based regulation of cell wall genes in controlling intrinsic organ size.

## Result

### Identification and Analysis of Putative CW-miRNAs in *Arabidopsis*

To identify CW-miRNAs in *A. thaliana*, we collected 572 genes annotated as cell wall biosynthesis related and 491 genes encoding proteins enriched in the Golgi apparatus (Parsons et al., 2012). Searching against the 427 annotated miRNAs in *A. thaliana*, coupling computational prediction with degradome sequencing analysis, we identified 23 putative CW-miRNAs that are predicted to target 78 genes pertinent to primary wall biosynthesis (Figure 1; Supplemental Table 1). Using 34 sequenced small RNA populations derived from six different organ types, we found that most of these miRNAs did not show strong organ specific expression pattern (Figure 1B). Together the CW-miRNAs account for 5.4% of all miRNAs annotated in *A. thaliana*. However, except miR156h that represses a gene encoding a pectin methylesterase inhibitor (Stief et al., 2014), miR773 that negatively regulates callose deposition in response to fungal infection (Salvador-Guirao et al., 2018), and miR827 that involves in phosphate homeostasis (Kant et al., 2011), functions of this cohort of miRNAs have not been investigated.

**Figure 1.**
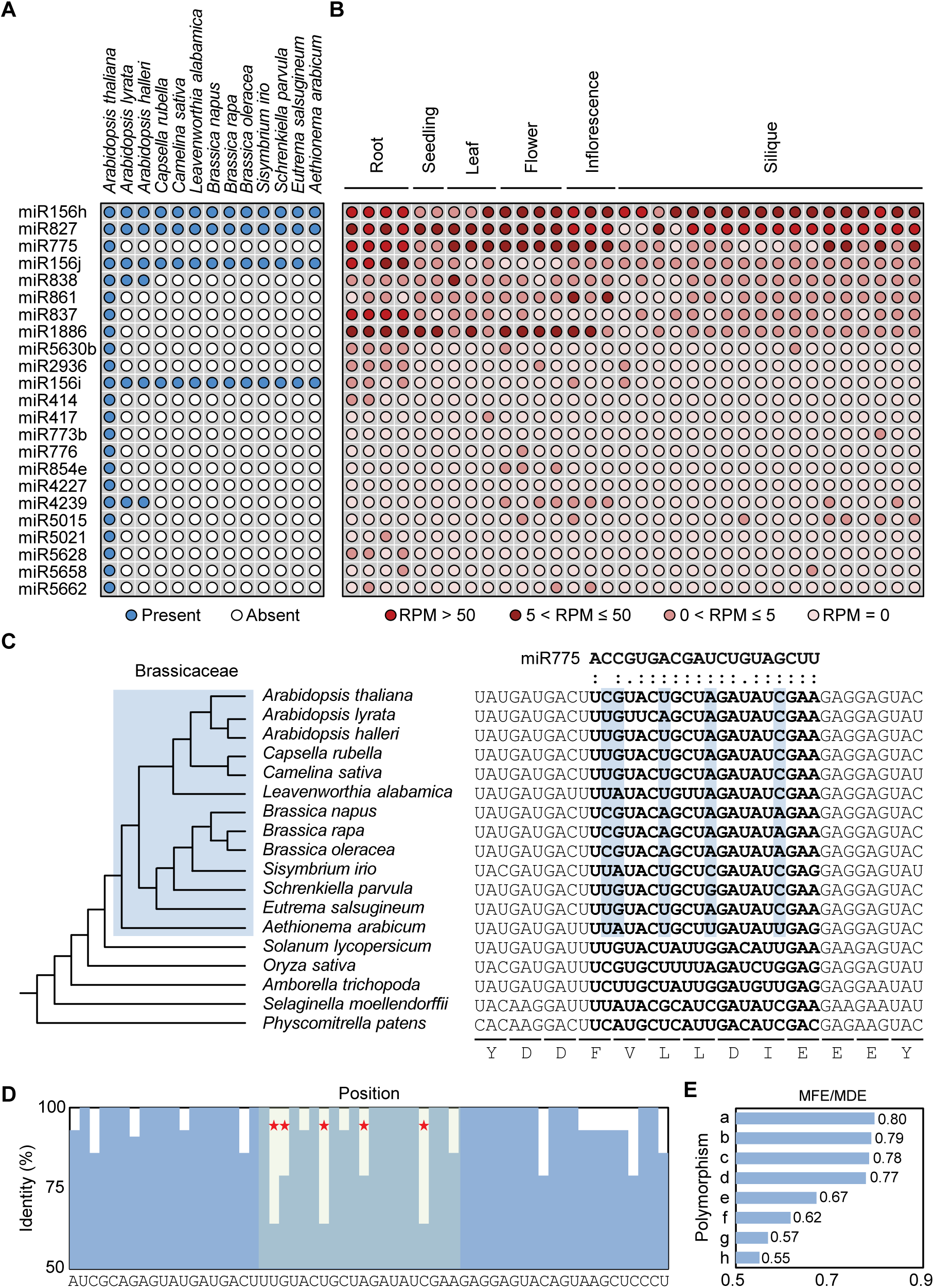
Identification and Analysis of Putative CW-miRNAs in *A. thaliana*. (**A**) Conservation of the 23 putative CW-miRNAs in *Brassicaceae*. Circles in blue indicate presence of a given CW-miRNA in the corresponding species. (**B**) Expression profile of the CW-miRNAs in *A. thaliana*. RPM (reads per million) values in 34 small RNA sequencing datasets, which are grouped into six organ types based on similarity of the sampled plant materials, are used to profile the miRNAs. (**C**) Comparison of the complementarity between miR775 and its possible binding site in *GALT9* homologs. On the left is a phylogenetic tree reconstructed with closest *GALT9* homologs from 18 species. Species in *Brassicaceae* are shaded in blue. On the right is an alignment of sequences flanking the miR775 binding site (in bold). The five polymorphic nucleotides within the miR775 binding site are shaded in green. (**D**) Quantification of nucleotide conservation in *GALT9* at the miR775 binding site across the 18 examined species. Red stars indicate the high-diversity nucleotides. The consensus sequence is shown below. (**E**) Calculated MFE/MED ratios for predicted miR775:target duplexes. Lower case letters represent observed combinations of the five polymorphic nucleotides. a, CGUAC; b, UGAAC; c, UGUAC; d, UGUGC; e, UAUAC; f, UAUCC; g, UAUUU; h, CGAAA.

Sequence comparison in representative *A. thaliana* ecotypes and 13 *Brassicaceae* species revealed that most (17 or 73.9%) CW-miRNAs are only found in *A. thaliana* (Figure 1A). For example, miR775 was among the first batch of non-conserved miRNAs annotated in *A. thaliana* (Rajagopalan et al., 2006). We found that miR775 is highly conserved in *A. thaliana* ecotypes but absent in *A. lyrata* and *A. halleri* (Supplemental Figure 1). Consistent with previous reports (Felippes et al., 2008), we found that the closest pre-miR775a homolog in *A. lyrata* misses the mature miR775 sequence (Supplemental Figure 1A) and could not fold into the stem-loop secondary structure typical for miRNA precursors (Supplemental Figure 1B). These results suggest that miR775 has evolved specifically in *A. thaliana* after its divergence from the common ancestor of the *Arabidopsis* genus.

On the other hand, 75 of the 78 (96.2%) predicted target genes for the CW-miRNAs have apparent orthologs in the Brassicaceae. *GALT9*, the predicted target gene for miR775, encodes a galactosyltransferase belonging to the carbohydrate-active glycosyltransferase 31 (Supplemental Figure 2). Sequence alignment revealed that the predicted miR775 binding site in *GALT9* contains five heterogeneous nucleotides across the examined *Brassicaceae* species (Figure 1C), more frequent than the surrounding sequences (Figure 1D). The five variable nucleotides have formed eight polymorphic combinations in the examined *Brassicaceae* species (Figure 1E). Among these and possible paralogs in *A. thaliana*, the miR775 binding site in *GALT9* exhibited the highest MFE/MED ratio (Supplemental Figure 2B), which is the ratio between the minimum free energy (MFE) of a predicted miRNA:target duplex to the minimum duplex free energy (MED) of the miRNA bound to a fully complementary sequence, an quantitative indicator for likelihood of miRNA targeting (Alves et al., 2009). These results indicate that complementarity of *GALT9* to miR775 was selected in *A. thaliana*.

### Molecular Validation of *GALT9* as a MiR775 Target

To validate *GALT9* as a miR775 target, we first performed the 5’ RNA ligation mediated-rapid amplification of cDNA ends (5’ RLM-RACE) assay (Llave et al., 2002). The detected 5’ ends of truncated *GALT9* transcript locate preferentially at the 14^th^ and 15^th^ nucleotides within the region complementary to miR775, counting from the 5’ end of miR775 (Figure 2). While this result supports miR775-guided *GALT9* cleavage, the detected transcript ends deviated by about four nucleotides from the conventional cleavage site between the 10^th^ and 11^th^ nucleotides of complementarity (Llave et al., 2002; German et al., 2009). We therefore performed degradome sequencing for further analysis. For comparison with the wild type, we generated miR775-overexpressing plants (*MIR775A-OX*) in which the enhanced Cauliflower Mosaic Virus *35S* promoter was used to drive pre-miR775a expression (Supplemental Figure 3). From the degradome sequencing data, we retrieved reads mapped to the predicted miR775 binding site in *GALT9*, which were enriched in *MIR775A-OX* relative to the wild type (Figure 2B). Closer inspection revealed that the enriched reads were not confined to a single nucleotide but concentrated in a region several nucleotides downstream of the 10^th^ position relative to the 5’ end of miR775 (Figure 2C). These results are consistent with the 5’ RLM-RACE data (Figure 2A) to support miR775-dependent cleavage of the *GALT9* transcript at unconventional sites.

**Figure 2.**
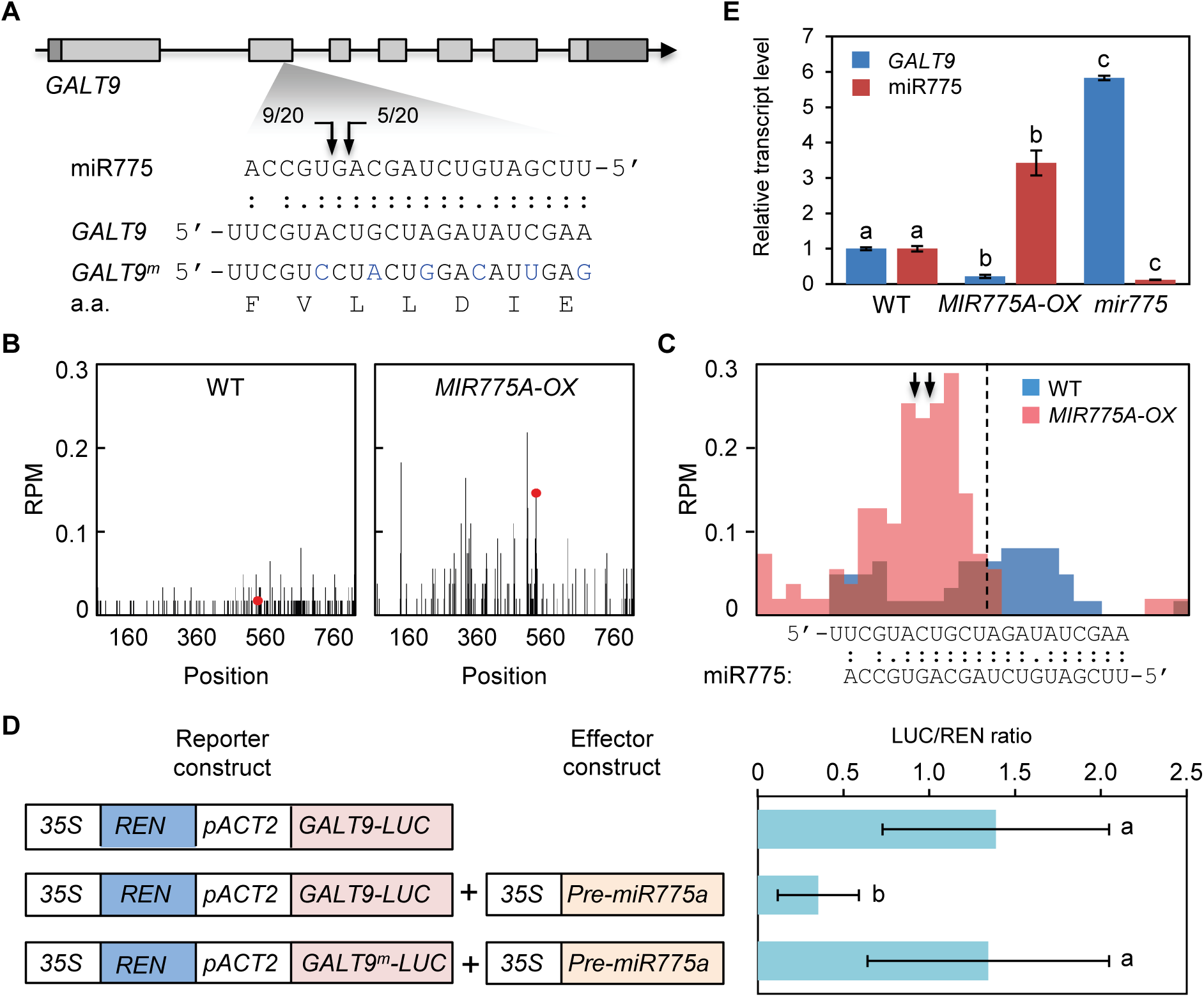
Validation of *GALT9* as an Authentic MiR775 Target. (**A**) 5’ RLM-RACE analysis of *GALT9*. Gene structure of *GALT9* is shown on top. Base pairing between miR775 and *GALT9* is shown on bottom. Arrows mark detected cleavage sites along with frequency of the corresponding clones. Substituted nucleotides for making *GALT9^m^* are colored in blue. (**B**) Comparison of degradome sequencing data obtained from the wild type (left) and *MIR775A-OX* (right) plants. Frequency of the sequenced 5’ ends is plotted against the position in the *GALT9* transcript. Red dots indicate position of reads with the highest frequency mapped to the miR775 binding site. (**C**) Sliding window analysis of degradome sequencing data at the miR775 binding site. Step of 4 nucleotides was used. Dashed line marks the position between the 10^th^ and 11^th^ nucleotides from the 5’ end of miR775. Arrows indicate positions of the cleavage sites mapped by 5’ RLM-RACE in A. (**D**) REN/LUC dual luciferase assay validating *GALT9* repression by miR775. The *Actin2* promoter was used to drive expression of *GALT9-LUC* or *GALT9^m^-LUC*. The *35S:pre-miR775a* effector and the reporters were used to transiently co-transform tobacco protoplasts. The LUC/REN ratio of chemiluminescence is shown on the right. Data are means ± SD from four independent transformation events. Different letters denote combinations with significant difference (Student’s *t*-test, *p* < 0.05). (**E**) Quantitative analysis of the miR775 and *GALT9* transcript levels in seedlings of the three indicated genotypes. Data are means ± SD from three technical replicates. Different letters denote groups with significant difference (Student’s *t*-test, *p* < 0.01).

Next, we tested whether miR775 is sufficient for repressing *GALT9* using the dual firefly luciferase (LUC) and *Renilla* luciferase (REN) reporter system (Liu et al., 2014). We generated a *GALT9-LUC* reporter construct in which the *GALT9* coding region was fused with that of *LUC* (Figure 2D). We also generated *GALT9^m^-LUC* by substituting the nucleotides of the miR775 binding site in *GALT9-LUC* but not the encoded amino acids (Figure 2A and 2D). Transient expression of these constructs in tobacco protoplasts showed that the LUC/REN chemiluminescence ratio was significantly lowered in the presence of miR775 (Figure 2D). Attenuation of the LUC/REN ratio was abolished in the *GALT9^m^-LUC* plus miR775 combination (Figure 2D), indicating that miR775 represses *GALT9-LUC* expression in a site-specific manner.

Finally, we examined how endogenous *GALT9* level is affected by genetic manipulation of miR775. In addition to the *MIR775A-OX* lines, we employed the CRISPR/Cas9 system to delete a 123 bp genomic region in *MIR775A* (the only locus in *A. thaliana*) encompassing miR775 (Supplemental Figure 4). Homozygous lines with no detectable expression of miR775 were selected and named *mir775* (Supplemental Figure 4B-4D). By quantitative reverse transcription coupled PCR (RT-qPCR) analysis, we found that the level of miR775 was significantly increased and decreased in *MIR775A-OX* and *mir775* in comparison to the wild type, respectively (Figure 2E). *GALT9* transcript level was significantly decreased in *MIR775A-OX* but increased in *mir775* relative to the wild type (Figure 2E). These results indicate that altering miR775 level is sufficient to reciprocally module *GALT9* transcript abundance.

### The *MIR775A*-*GALT9* Circuit Controls Organ and Cell Sizes

To elucidate the biological role of miR775, we monitored morphology of the *mir775* plants throughout the life cycle. In comparison to the wild type, a size reduction of leaf-related organs, including the cotyledon, the fifth rosette leaf, and the petal, was observed for *mir775* (Figure 3; Supplemental Figure 4E-4H). Quantification confirmed that *mir775* has significantly smaller phyllome organs than the wild type (Figure 3D-3F). In contrast, mature organs of *MIR775A-OX* were significantly larger than those of the wild type (Figure 3). To confirm the *mir775* phenotype, we generated the *MIR775A-OX mir775* double mutant through genetic crossing (Supplemental Figure 5). We found that the *35S:pre-miR775a* transgene in the used *MIR775A-OX* line was able to restore miR775 transcript accumulation and rescue the organ reduction phenotype in the *mir775* background (Figure 3; Supplemental Figure 5).

**Figure 3.**
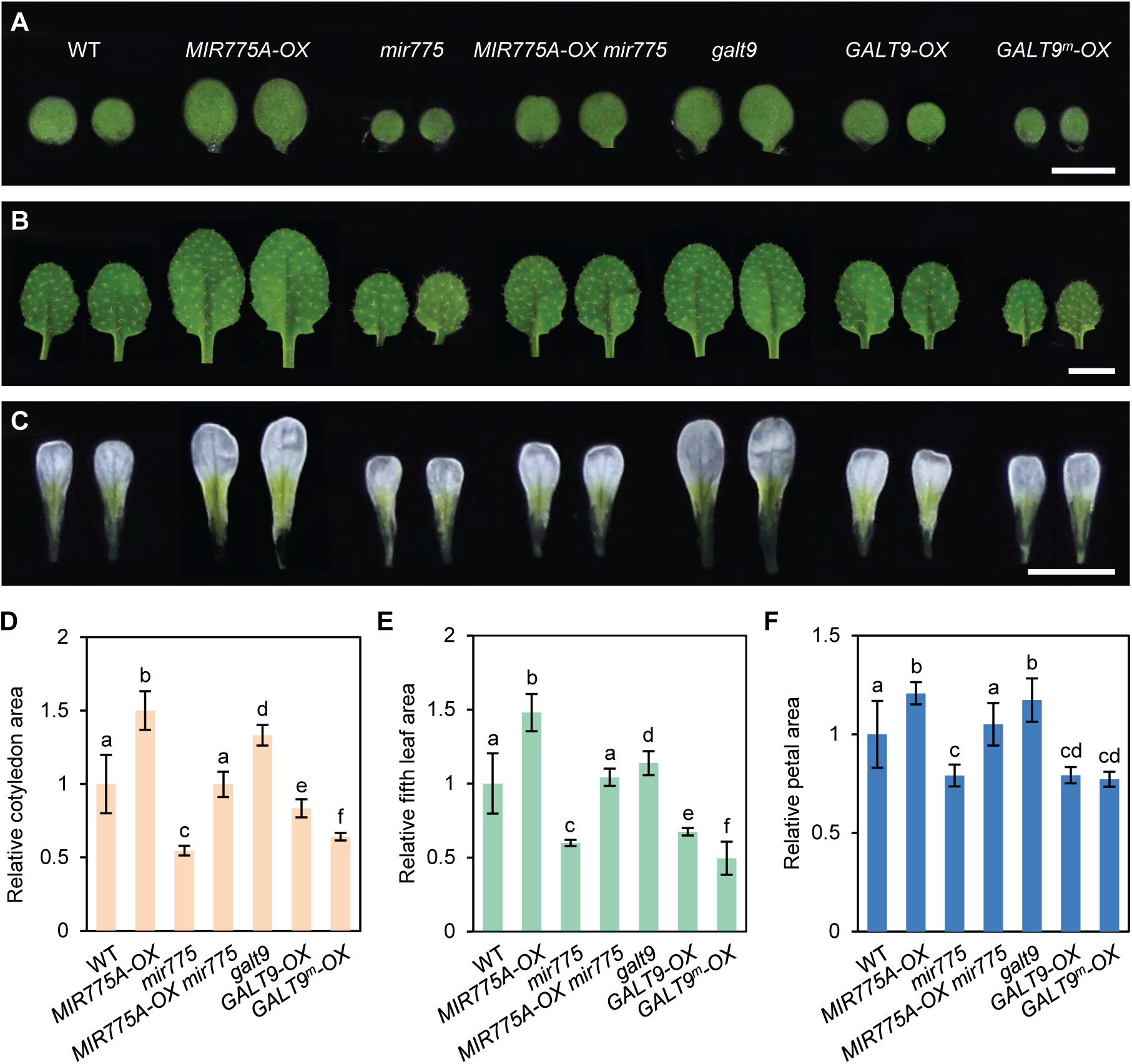
*MIR775A* and *GALT9* Oppositely Regulate Size of Leaf-related Organs. (**A-C**) Morphological comparison of three representative organ types across the indicated genotypes. (A) Cotyledon of seven-day-old seedlings; (B) The fifth rosette leaf of three-week-old plants; (C) petal of open flowers. Bars, 2 mm. (**D-F**) Quantitative size measurement of cotyledons (D), the fifth rosette leaves (E), and the petals (F). Data are mean ± SD from individual organs normalized against the wild type. Different letters denote genotypes with significant difference (Student’s *t*-test, n = 30, *p* < 0.001 for D, n = 20, *p* < 0.01 for E, n = 30, *p* < 0.001 for F).

**Figure 4.**
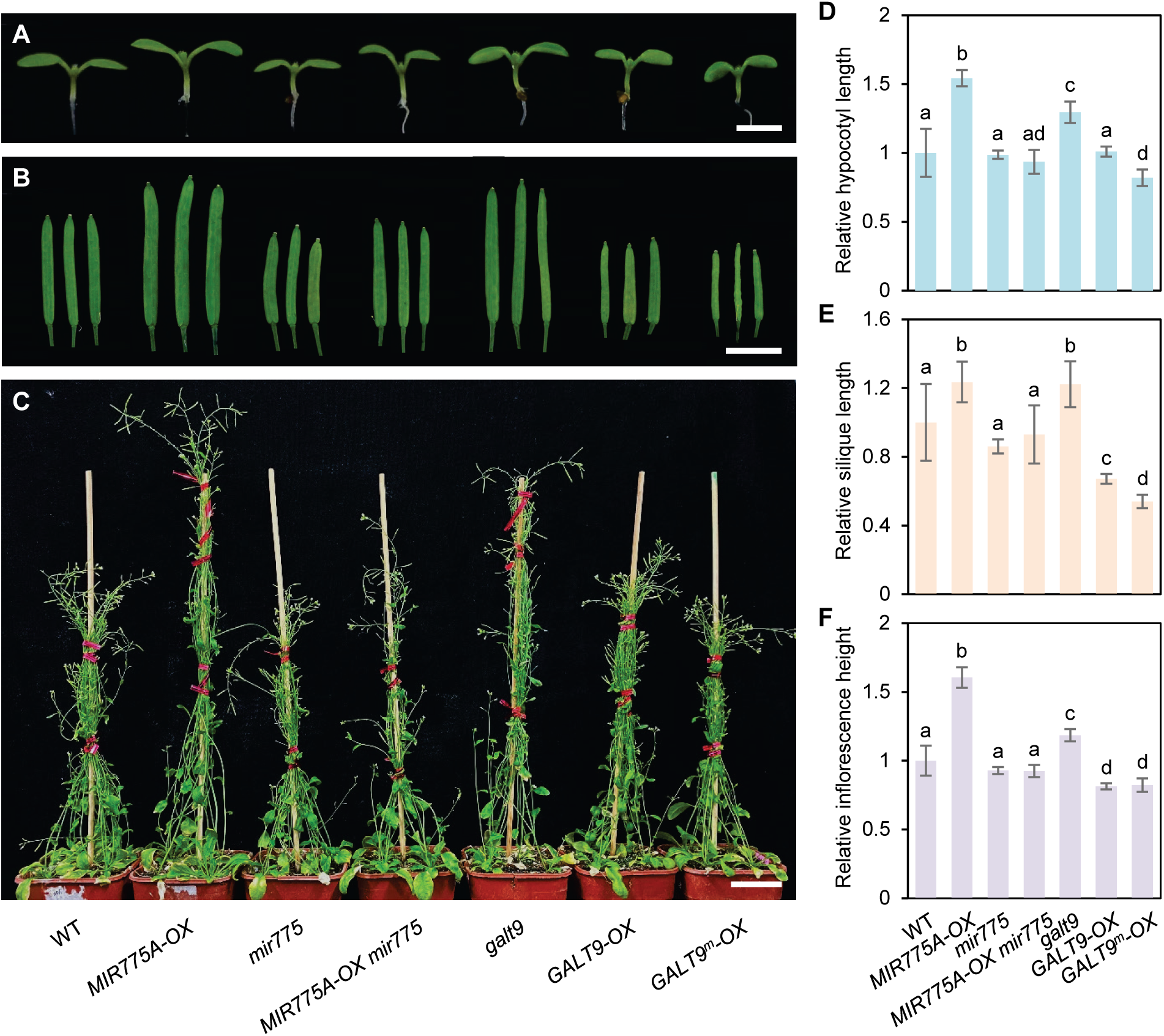
*MIR775A* and *GALT9* Play Different Roles in Regulating Size of Heterotrophic Organs. (**A-C**) Morphological comparison of three representative organs with heterotrophic growth across the indicated genotypes. (A) Hypocotyl of seven-day-old seedlings, bar, 2 mm; (B) Mature silique, bar, 2 mm; (C) Mature inflorescence, bar, 2 cm. (**D-F**) Quantitative measurement of hypocotyl length (D), silique length (E), and inflorescence height (F). Values are mean ± SD from individual organs normalized to the wild type. Different letters denote genotypes with significant difference (Student’s *t*-test, n = 15, *p* < 0.01 for D, n = 30, *p* < 0.001 for E, n = 26, *p* < 0.001 for F).

To test the role of *GALT9* in phyllome organs, we employed the CRISPR/Cas9 system to delete the entire coding region of *GALT9* (Supplemental Figure 6). In the homozygous deletion lines (*galt9-1*), *GALT9* expression was drastically compromised in comparison with the wild type (Supplemental Figure 6A-6C). We also identified an *Arabidopsis* T-DNA line (*galt9-2*) carrying insertion in the start codon of *GALT9* (Supplemental Figure 6A). Both *galt9* mutants exhibited significantly enlarged phyllome organs than the wild type (Figure 3), phenotypes similar to *MIR775A-OX*. We also generated transgenic plants over-expressing *GALT9* (*GALT9-OX*) and *GALT9^m^* (*GALT9^m^-OX*; see Figure 2A) driven by the *35S* promoter (Supplemental Figure 7). Both *GALT9-OX* and *GALT9^m^-OX* plants displayed significantly reduced sizes of leaf-related organs than the wild type (Figure 3; Supplemental Figure 7), phenotypes similar to those of *mir775*.

In contrast to the phyllome, there are organs in *A. thaliana* that rely on heterotropic growth to reach the intrinsic sizes, such as the hypocotyl, the silique, and the inflorescence stem (Geitmann and Ortega, 2009; Peaucelle et al., 2015; Andres-Robin et al., 2018). In comparison to the wild type, we found that hypocotyl length, silique length, and inflorescence height of the *mir775* plants were not statistically different from those of the wild type (Figure 4). By contrast, sizes of these organs of the *MIR775A-OX*, *galt9*, *GALT9-OX*, and *GALT9^m^-OX* plants were significantly altered compared to the wild type with the exception of hypocotyl length of *GALT9-OX* (Figure 4). Collectively, these results indicate that endogenous miR775 primarily promotes phyllome organ growth by repressing *GALT9* in *A. thaliana*.

In addition to *GALT9*, we have previously reported three other computationally predicted target genes for miR775 including *DICER-LIKE1* (*DCL1*) (Zhang et al., 2011). Inspection of the degradome sequencing data from both the wild type and *MIR775A-OX* backgrounds revealed no evidence for miR775-directed cleavage for these genes (Supplemental Figure 8). Furthermore, consistent with previous characterizations of the *dcl1* mutants (e.g. Mallory and Vaucheret, 2006), an examined *dcl1* T-DNA insertion mutant exhibited significantly reduced organ sizes in comparison to the wild type (Supplemental Figure 9), phenotype opposite to that of *galt9* or *MIR775A-OX*. Thus, *GALT9* is a *bona fide* miR775 target that plays an opposite role to miR775 in determining intrinsic organ size.

A change in organ size can be attributed to altered cell size and/or cell number. To assess the effects of the *MIR775A-GALT9* circuit, we selected four cell types from three organs for examination by cryo-scanning electron microscopy (cryo-SEM). Observed and quantified sizes of *MIR775A-OX* and *galt9* epidermal cells on the cotyledon, the petal, and the hypocotyl as well as the guard cells on the cotyledon were significantly larger than those of the wild type (Figure 5). Opposite phenotypes were observed for *mir775* and *GALT9-OX* cells (Figure 5A-5E). Moreover, a highly linear relationship with a virtually 1:1 slope between the cell size and the organ size was observed for the three examined organ types across the five genotypes (Figure 5F). These results indicate that changes in cell size are primarily responsible for changes in organ size caused by manipulating the *MIR775A-GALT9* circuit.

**Figure 5.**
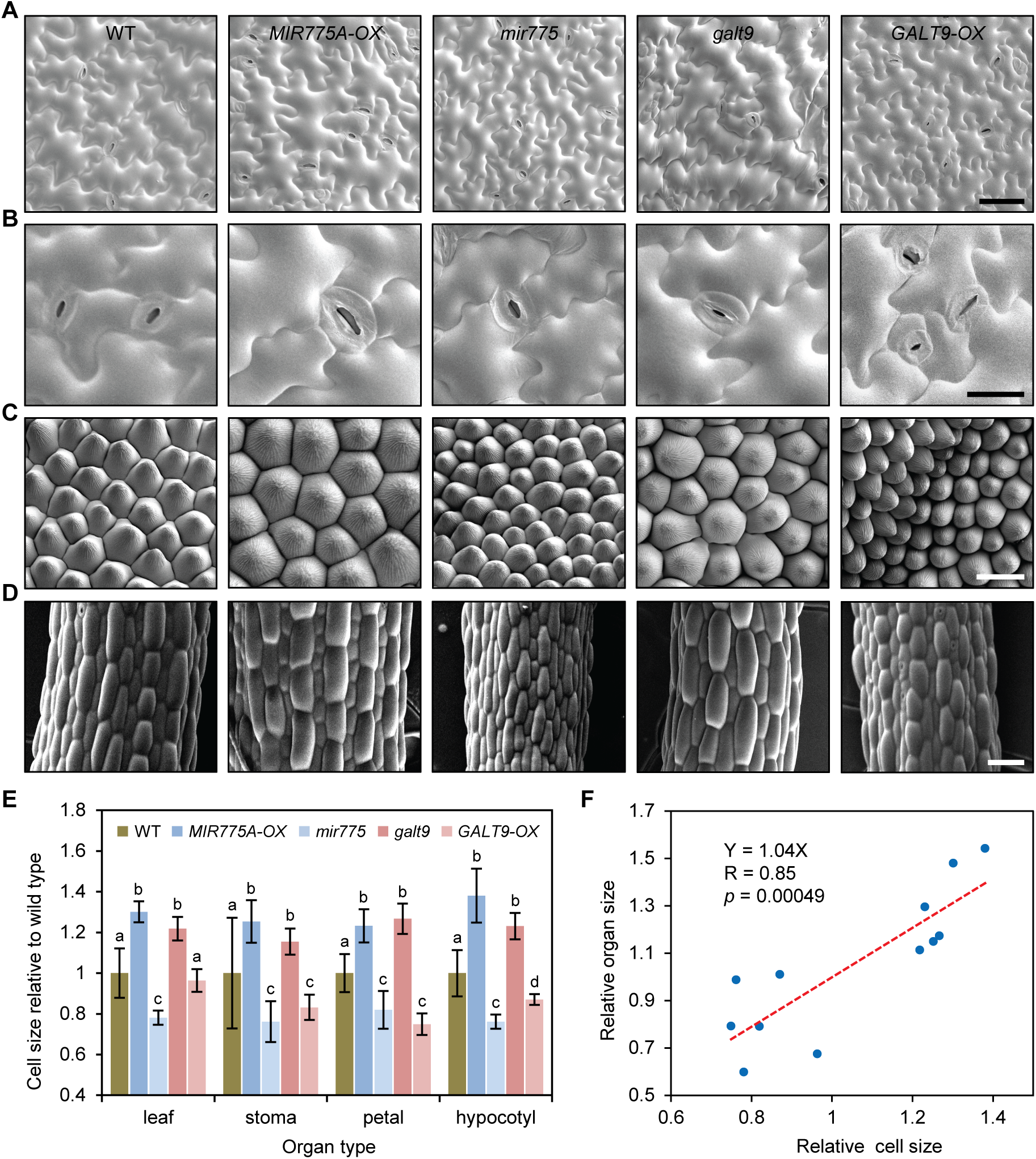
The *MIR775A-GALT9* Circuit Controls Cell Size. (**A-D**) cryo-SEM analysis of epidermal cells of the five indicated genotypes. Shown are representative images for cotyledon (A), bar, 50 μm; stoma including guard cells (B), bar, 20 μm; petal (C), bar, 20 μm; and hypocotyl (D), bar, 50 μm. (**E**) Quantification of epidermal cell size from cotyledon, petal, and hypocotyl and stoma area. Data are mean ± SD relative to the wild type from 30 individual cells of several individual plants. Different letters denote genotypes with significant difference (Student’s *t*-test, *p* < 0.01 for A, C and D, *p* < 0.05 for B). (**F**) Correlation between cell size and organ size. Relative organ and cell sizes of three organs (cotyledon, petal, and hypocotyl) across the wild type, *MIR775A-OX*, *mir775*, *galt9*, and *GALT9-OX* genotypes were used for a linear regression analysis.

**Figure 6.**
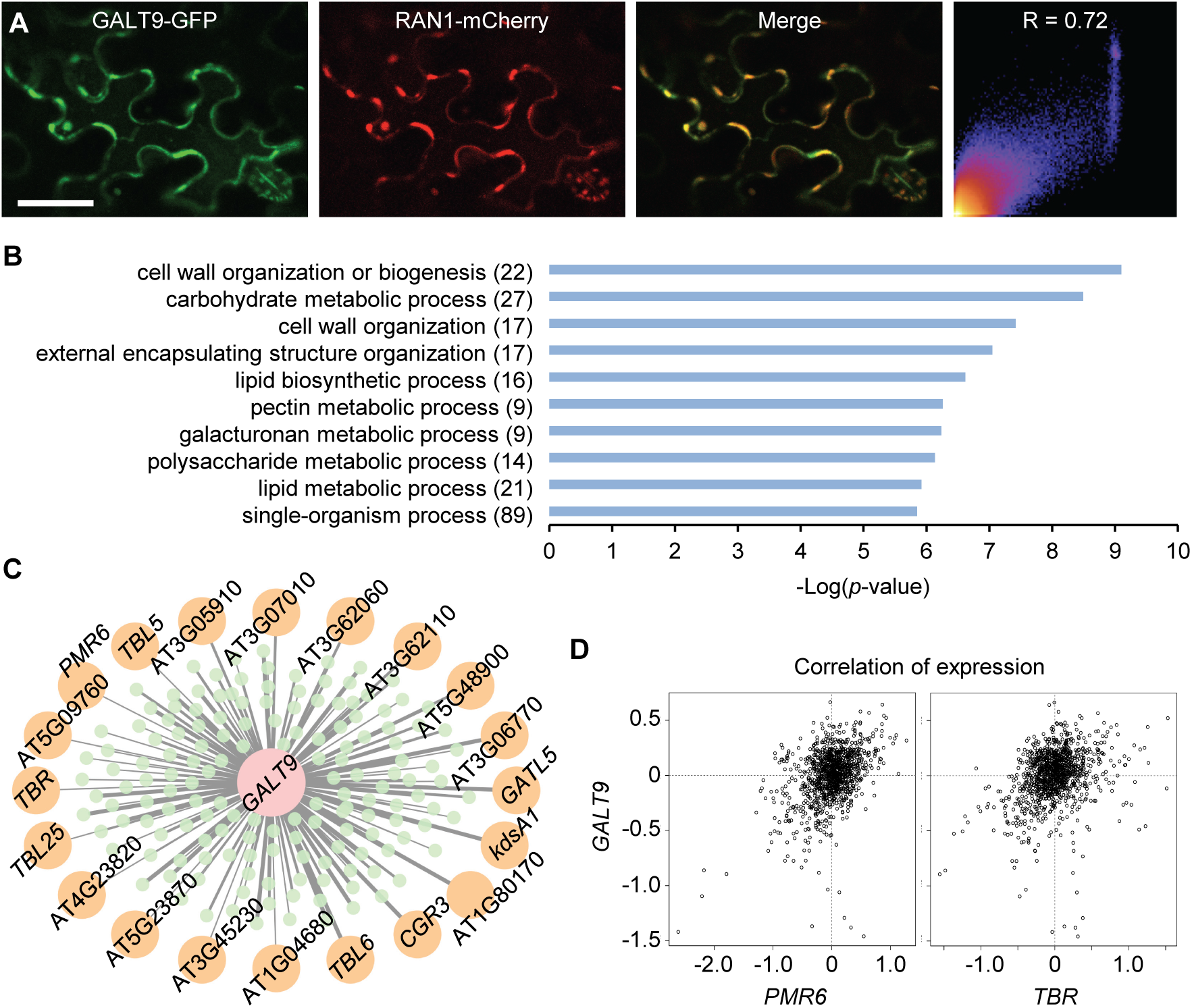
*GALT9* Has a Deduced Role in Pectin Metabolism. (**A**) Colocalization of GALT9-GFP with RAN1-mCherry in tobacco leaf epidermal cells. Scatter plot on the right shows correlation of GFP and mCherry fluorescence intensity. R, Pearson correlation coefficient. Bar, 50 μm. (**B**) Top ten most significantly enriched GO terms in the biological process category associated with the 174 *GALT9* co-expressed genes. Numbers in parentheses are co-expression genes associated with each term. (**C**) Concentric display of *GALT9* co-expression genes with the 20 pectin-related genes shown on the periphery. Narrow lines representing mutual rank value above 200, medium lines representing 50-200, and wide lines representing 0-50. (**D**) Correlation pattern between *GALT9* and the pectin-related genes *PMR6* and *TBR*. Axes are Log_2_-transformed expression levels against the averaged level of each gene.

### *MIR775A*-*GALT9* Modulates Pectin Level and Cell Wall Elasticity

Members of the *GALT* family have been extensively implicated in cell wall remodeling (Supplemental Figure 2A) (Bouton et al., 2002; Qu et al., 2008; Qin et al., 2017). As most proteins involved in cell wall remodeling locate on the endomembrane (Parsons et al., 2012), we determined the subcellular localization of GALT9. RESPONSIVE TO ANTAGONIST1 (RAN1) is a copper transporter reported to reside on the endomembrane (Hirayama et al., 1999). Using GALT9 fused with the green fluorescent protein (GFP), we found that GALT9-GFP colocalized with mCherry-tagged RAN1 transiently co-expressed in the same tobacco leaf epidermal cells (Figure 6). This observation indicates that transiently expressed GALT9 is located on the endomembrane.

To infer the molecular function of *GALT9*, we carried out a co-expression analysis and identified 174 genes that are co-expressed with *GALT9* in *A. thaliana* (Supplemental Dataset 1). Gene Ontology (GO) analysis revealed that these genes were most significantly enriched with GO terms related to cell wall biology and pectin metabolism in particular (Figure 6B). Manual review revealed that 20 of these genes are linked to pectin metabolism and related processes, including eight genes of the pectin lyase-like superfamily, four genes of the *TRICHOME BIREFRINGENCE-LIKE* family, and eight other genes in pectin synthesis and modifications based on experimental evidence in the literature (Figure 6C). As examples, co-expression patterns between *GALT9* and *TRICHOME BIREFRINGENCE* (*TBR*), which was shown to regulate pectin composition in the trichome and stem (Bischoff et al., 2010), and between *GALT9* and *POWDERY MILDEW RESISTANT6* (*PMR6*), a member of the pectin lyase-like superfamily and whose mutation caused smaller rosette leaves with altered pectin composition (Vogel et al., 2002), are shown in Figure 6D.

To confirm the involvement of GALT9 in pectin metabolism, we performed monosaccharide composition analysis of the cell walls. We found that the relative amount of glucose, the primary monosaccharide of cellulose, was not significantly different in the de-starched fifth rosette leaves from the *mir775*, *MIR775A-OX*, *galt9*, and *GALT9-OX* plants in comparison to the wild type (Figure 7). In contrast, the relative amount of galacturonic acid, the representative derivative of pectin polysaccharides, was significantly lower in the *MIR775A-OX* and *galt9* plants but higher in the *mir775* and *GALT9-OX* plants than the wild type (Figure 7A). Moreover, an inverse linear relationship between the relative amount of galacturonic acid and the relative cell size was observed among the five genotypes (Figure 7B). This linear relationship was not found for the relative glucose level (Figure 7B). These results indicate that *MIR775A-GALT9* specifically influences pectin level in the leaf cell walls.

**Figure 7.**
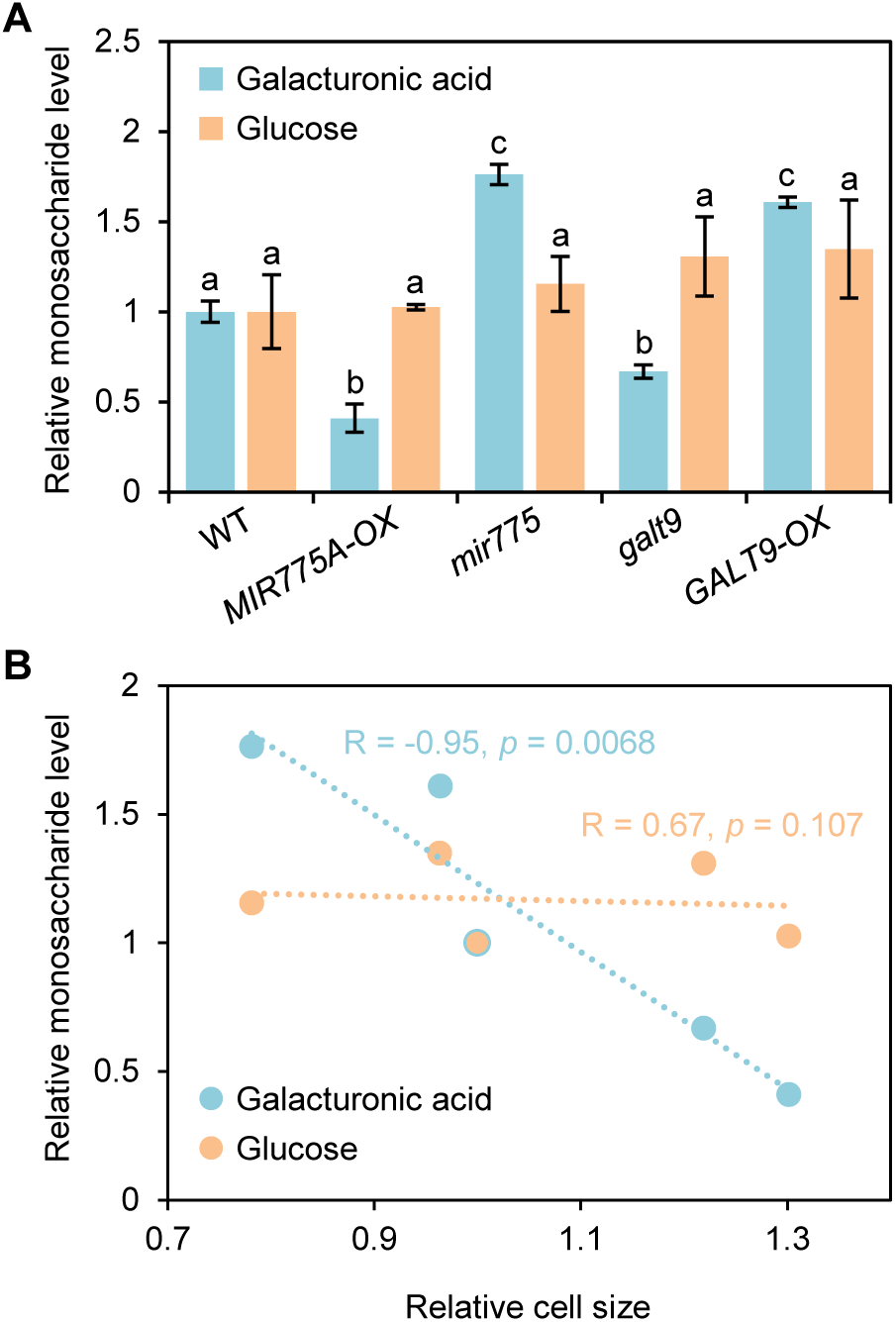
The *MIR775A-GALT9* Circuit Regulates Cell Wall Pectin Level. (**A**) Quantification of the relative glucose and galacturonic acid levels in the cell walls. Hydroxylated cell wall materials extracted from the de-starched fifth rosette leaf of the indicated genotypes were used for monosaccharide measurement by colorimetry. Data are mean ± SD from three technical replicates performed on pooled leaves. Within a set of measurements, different letters denote genotypes with significant difference (Student’s *t*-test, *p* < 0.01). (**B**) Correlation between relative cell size and the two quantified cell wall monosaccharides across the five genotypes by a linear regression analysis.

Raman imaging is a technique for obtaining high-resolution, chemically specific, and non-destructive information of plant cell walls (Gierlinger et al., 2012; Zeng et al., 2016). Using a home-built coherent Raman microscope, we mapped *in situ* pectin distribution in a mutant defective in *QUARTET2* (*QRT2*). Stronger than wild type signals encircling cotyledon epidermal cells were observed in *qrt2* (Supplemental Figure 10), consistent with previous reports that QRT2 is required for pectin degradation (Rhee and Somerville, 1998). Similar to *qrt2*, we detected stronger than wild type pectin signals in both *mir775* and *GALT9-OX* plants (Supplemental Figure 10A). The *MIR775A-OX* and *galt9* plants, in contrast, exhibited the opposite phenotype with weaker pectin signals than the wild type (Figure 8). This effect was specific for pectin, as no difference in cellulose deposition among *MIR775A-OX*, *galt9*, and the wild type was observed (Figure 8A and 8C). Quantification of the signal intensity confirmed that pectin content was significantly reduced in *MIR775A-OX* and *galt9* (Figure 8D).

**Figure 8.**
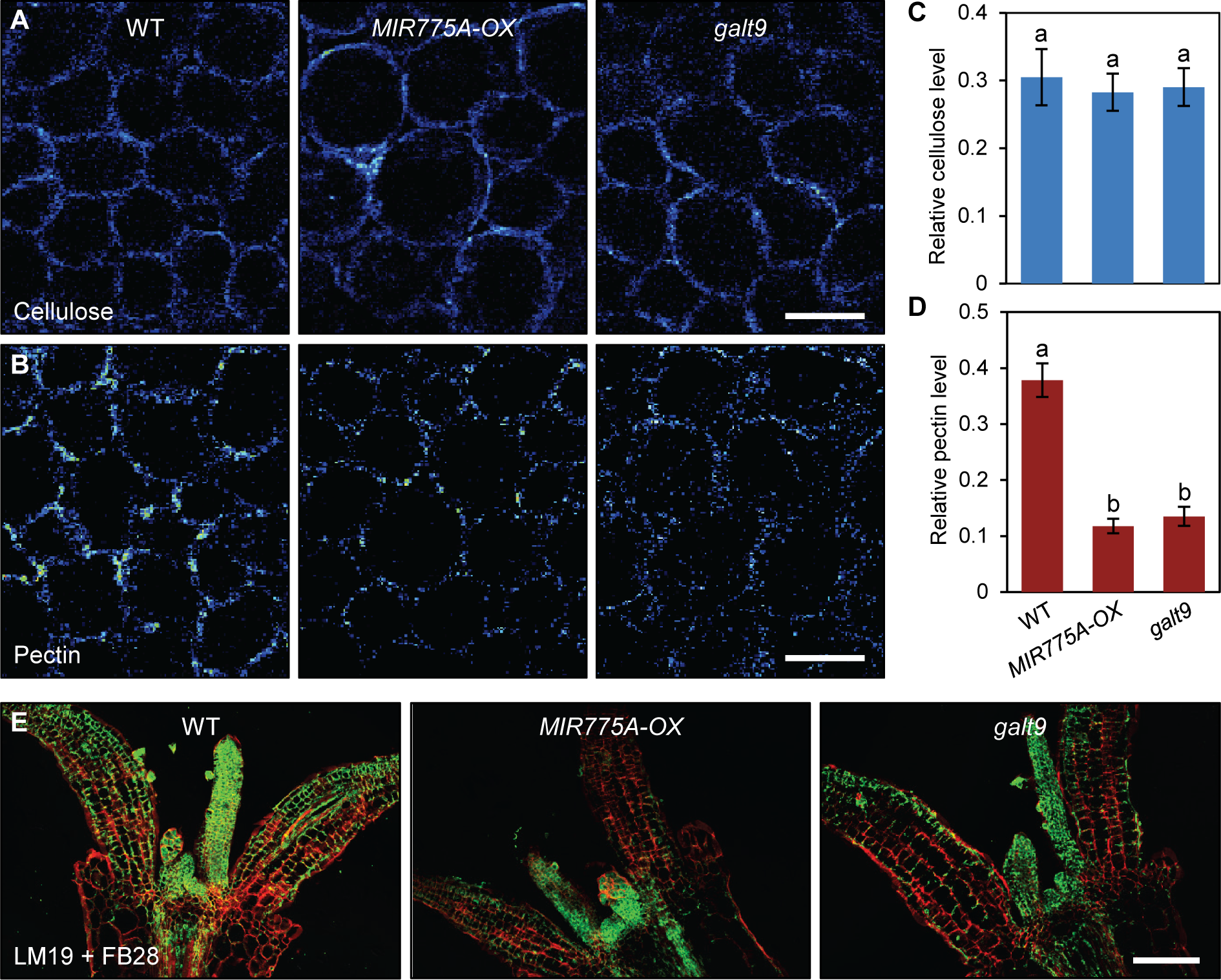
*MIR775A-OX* and *galt9* Seedlings Have Reduced Cell Wall Pectin. (**A-B**) Examination of cell wall constituents by confocal Raman microscopy. Cotyledon mesophyll cells of seven-day-old wild type, *MIR775A-OX*, and *galt9* seedlings were imaged for cellulose (A) at 1100 cm^−1^ and pectin (B) at 854 cm^−1^. Bars, 50 μm. (**C-D**) Relative cellulose and pectin levels deduced from Raman images. Average intensity in a 25 μm by 25 μm area at the cell corner was used to represent the level of the wall components. Data are mean ± SD of 15 areas from five cotyledons. Different letters denote genotypes with significant difference (Student’s *t*-test, *p* < 0.01). (**E**) Immunohistochemical localization of pectin. The LM19 antibody (green) and the FB28 dye (red) were used to stain seven-day-old seedlings and examined by fluorescence microscopy. Bar, 100 μm.

As HG accounts for more than 60% of plant cell wall pectin (Caffall and Mohnen, 2009), we performed immunohistochemical analysis of cotyledons using a fluorescence-labeled monoclonal antibody (LM19) specific for HG (Verhertbruggen et al., 2009). Fluorescence microscopy revealed that LM19 signals in the *MIR775A-OX* and *galt9* seedlings were drastically reduced in comparison to the wild type (Figure 8E). By contrast, Fluorescent Brightener 28 (FB28), which mainly stains cellulose, generated signals with no obvious difference among the genotypes (Figure 8E). These results confirmed that miR775 and GALT9 reduces and promotes pectin deposition in the cell walls, respectively.

AFM is useful for determining the surface structures and mechanical characters of biological samples at the nanometer scale (Yakubov et al., 2016). To investigate the link between pectin content and mechanical property of the cell wall, we employed AFM to directly measure the elastic properties of the epidermal cells. This analysis showed that the *qrt2* mutant has higher elastic modulus than the wild type (Supplemental Figure 10B), consistent with the notion that higher pectin level leads to increased stiffness of the wall. We then applied AFM to measure the elastic properties of the *MIR775A-OX* and *galt9* cotyledon cells and petal cells (Figure 9). In accordance with the cryo-SEM results (Figure 5), the 3D contour mapped by AFM revealed that the *MIR775A-OX* and *galt9* cells are larger than the wild type (Figure 9A and 9D). The *MIR775A-OX* and *galt9* cell walls, however, have elastic moduli significantly lower than the wild type (Figure 9C and 9F), indicating that the enlarged cells have reduced wall rigidity. Taken together, our results demonstrate that *MIR775A-GALT9* modulates pectin abundance in the cell wall and affects resistance to micro-indentation.

**Figure 9.**
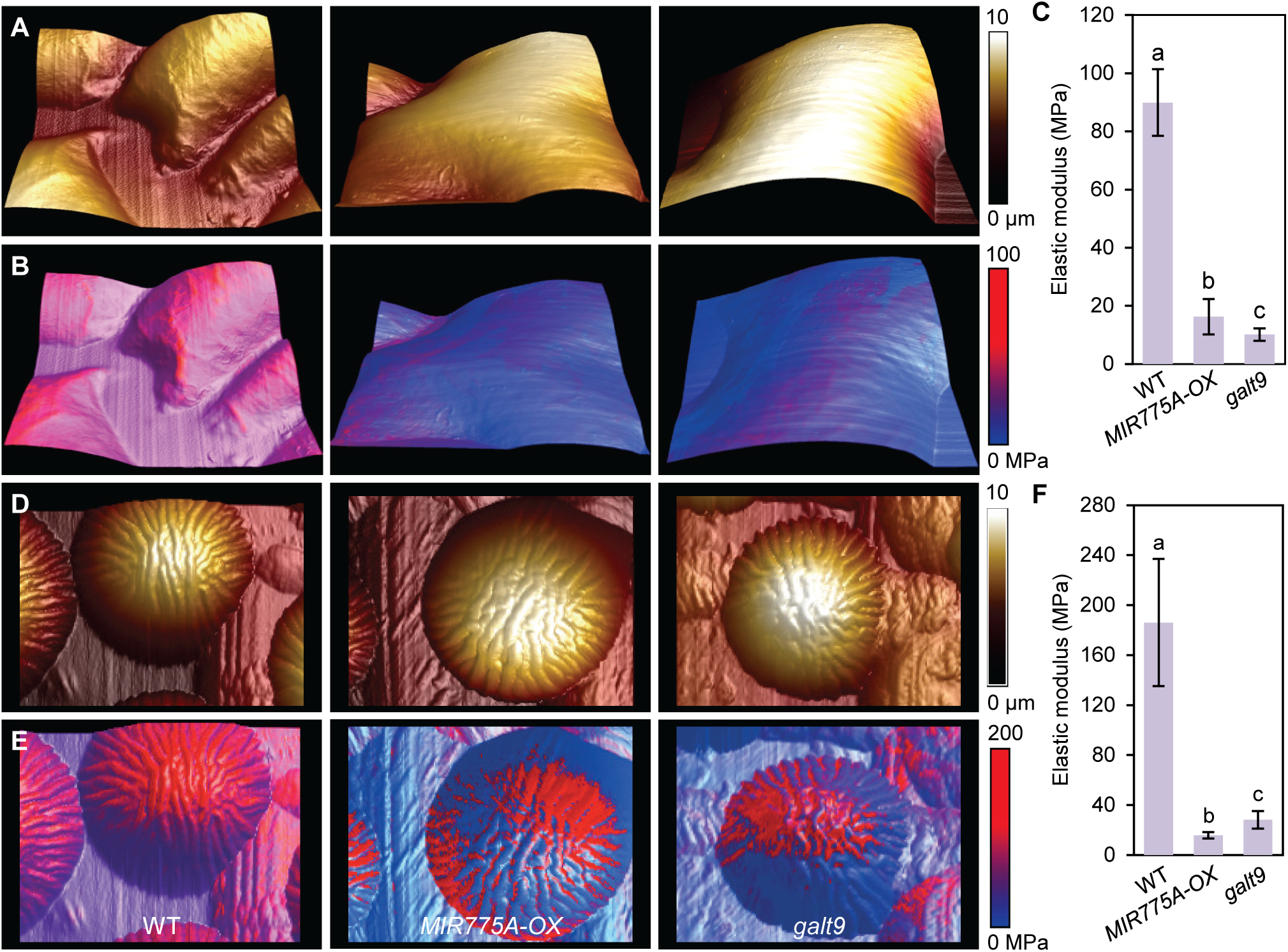
*MIR775A-OX* and *galt9* Epidermal Cells Have Reduced Elastic Modulus. (**A**) AFM mapping of three-dimensional topography of epidermal cells. Individual cells of seven-day-old cotyledons were analyzed. Colors represent distance from the base, which is the deepest point the probe reaches. (**B**) Cell topography overlaid with elastic modulus. Colors indicate elasticity. (**C**) Quantification of apparent Young’s modulus using the Peak Force QNM mode. Each measurement was the average in a 5 μm by 5 μm area of a cell with the highest modulus. Data are mean ± SD of 10 cells from three cotyledons. Different letters denote genotypes with significant difference (Student’s *t*-test, *p* < 0.001). (**D-F**) Cell topography (D), topography overlaid with elasticity (E), and apparent Young’s modulus (F) of the petal epidermal cells. Individual cells of petals of open flowers were analyzed. Each measurement was the average in a 10 μm by 10 μm area with the highest modulus. Data are mean ± SD of 10 cells from three petals. Different letters denote significant difference (Student’s *t*-test, *p* < 0.001).

### *MIR775A* Is Negatively Regulated by *HY5* in Aerial Organs

A full-length cDNA *BX81802* matches the *MIR775A* locus, allowing the transcription start site and proximal promoter region (*pMIR775A*) to be determined (Figure 10). To find out how *MIR775A* is transcriptionally regulated, we examined available whole genome chromatin immunoprecipitation (ChIP) data and identified an ELONGATED HYPOCOTYL5 (HY5) binding peak in *pMIR775A* (Figure 10A) (Zhang et al., 2011). As a key transcription factor for photomorphogenesis, HY5 is known to bind to G-box-like motifs (Oyama et al., 1997; Yadav et al., 2002; Song et al., 2008). Indeed, we located a typical G-box like motif in *pMIR775A* that coincides with the HY5 binding peak (Figure 10A). Using ChIP-qPCR, significant enrichment of HY5 occupancy at *pMIR775A* was confirmed (Figure 10B). These results reveal *HY5* as a plausible upstream regulator for the *MIR775A-GALT9* circuit.

**Figure 10.**
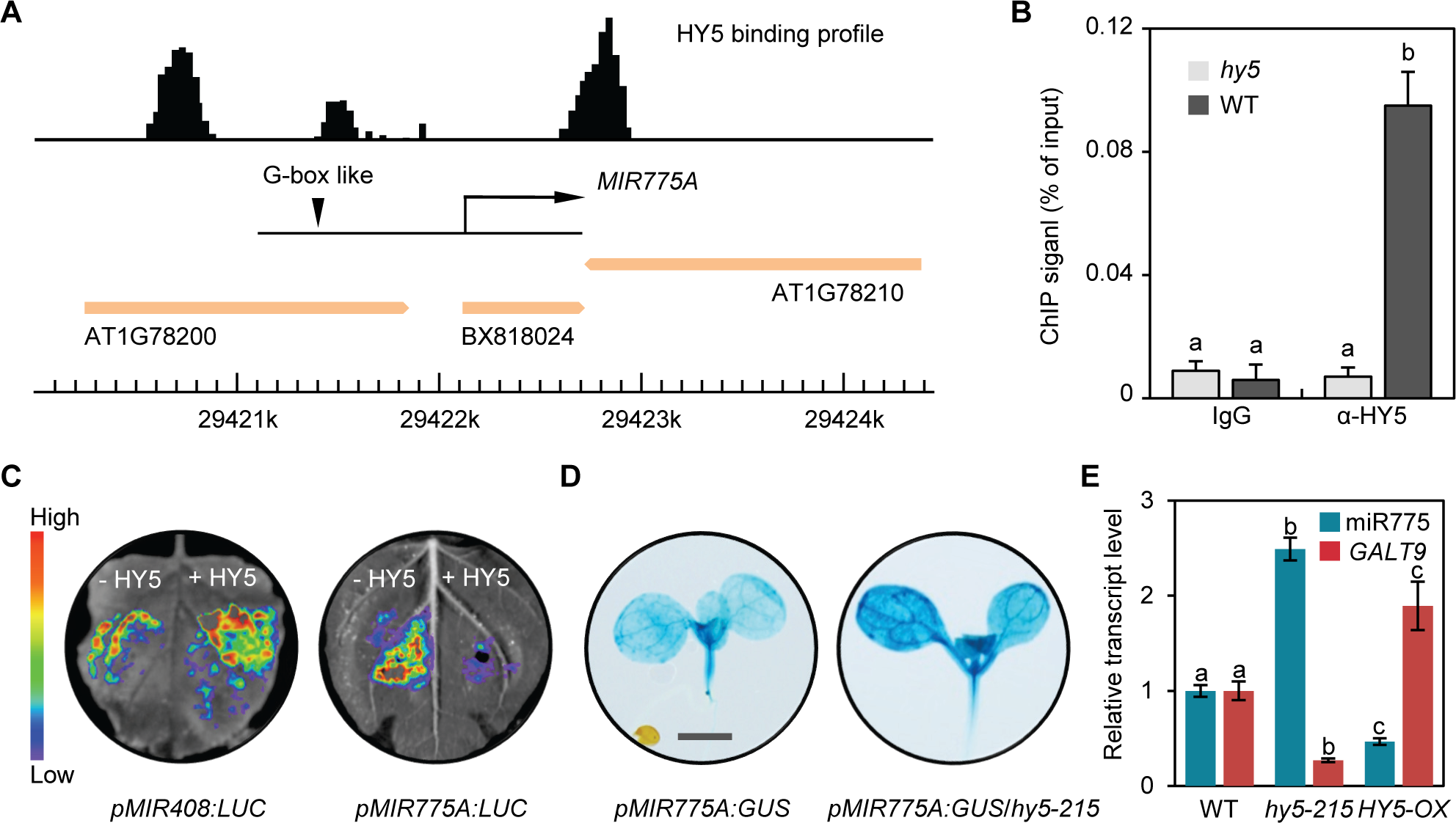
HY5 Represses *MIR775A* Expression by Directly Binding to Its Promoter. (**A**) HY5 occupancy profile at the *MIR775A* locus. HY5 binding profile is based on global ChIP data mapped onto the *Arabidopsis* genome coordinates. Loci are represented by block arrows. Position of *MIR775A*, defined by the full-length cDNA *BX818024*, is depicted as a black arrow. The triangle marks the G-box like motif. (**B**) Confirmation of HY5 binding to *pMIR775A* by ChIP-qPCR. ChIP was performed in light-grown wild type and *hy5* seedlings with or without the anti-HY5 antibody. Values are normalized to the respective DNA inputs. Data are ± SD from three technical replicates. Different letters denote significant difference (Student’s *t*-test, *p* < 0.001). (**C**) Transient expression assay for testing the effect of HY5 on *pMIR775A* activity. Either the *pMIR775A:LUC* or *pMIR408:LUC* construct was co-infiltrated with the *35S:HY5-GFP* (+HY5) or the vector alone (-HY5) in tobacco epidermal cells and imaged for LUC activity. (**D**) GUS staining for *HY5*-dependent *pMIR775A* activity in *A. thaliana*. The same *pMIR775A:GUS* reporter gene was expressed in either the wild type or the *hy5-215* background. Bar, 1 mm. (**E**) RT-qPCR analysis of the relative miR775 and *GALT9* transcript abundance in the wild type, *hy5-215*, and *HY5-OX* seedlings. Data are means ± SD from three technical replicates. Different letters denote groups with significant difference (Student’s *t*-test, *p* < 0.01).

To examine the effect of HY5 on *pMIR775A in vivo*, we generated the *35S:GFP* and *35S:HY5-GFP* effector constructs. As reporters, we used *pMIR775A* to drive *LUC* and *pMIR408*, which was previously shown to be activated by HY5 (Zhang et al., 2014), as a positive control. We tested four effector-reporter combinations through co-infiltration of tobacco leaf epidermal cells. Attesting to validity of the assay, co-expression of HY5 with *pMIR408:LUC* robustly enhanced LUC activity (Figure 10C). However, in the presence of HY5, the *pMIR775A* activity was markedly decreased (Figure 10C), indicating that HY5 negatively regulates *MIR775A*. To corroborate this regulatory relationship in *A. thaliana*, we fused the *β-glucuronidase* (*GUS*) gene with *pMIR775A* and expressed the reporter in either the wild type (*pMIR775A:GUS*) or the *hy5-215* (*pMIR775A:GUS*/*hy5-215*) genetic background (Figure 10D). In both seedlings and adult plants, we found that GUS activity in the shoot was higher in *hy5-215* than in the wild type (Figure 10D; Supplemental Figure 11), confirming *HY5*-mediated *MIR775A* repression.

Finally, we performed RT-qPCR analysis to monitor the influence of *HY5* on miR775 and *GALT9* transcript accumulation. For this purpose, we also employed a *HY5-OX* line in which expression of the *HY5* coding region was driven by the *35S* promoter (Gao et al., 2020). This analysis revealed that miR775 abundance increased in the *hy5-215* shoots but decreased in *HY5-OX* with reference to the wild type (Figure 10E). Conversely, *GALT9* transcript level was significantly lower in *hy5-215* but higher in *HY5-OX* shoots compared to the wild type (Figure 10E). Collectively these results indicate that HY5 binds to the *MIR775A* promoter to repress miR775 accumulation and derepress *GALT9* in aerial organs, thereby forming the *HY5-MIR775A-GALT9* repression cascade.

Previously, we reported that *HY5* positively regulates *MIR775A* based on analysis of miR775 abundance in whole young seedlings (Zhang et al., 2011). To ascertain whether HY5 positively or negatively regulates *MIR775A*, we compared GUS activities in different organs of *pMIR775A:GUS* and *pMIR775A:GUS*/*hy5-215* plants. This analysis revealed that, in contrast to the aerial organs, GUS activity in *pMIR775A:GUS*/*hy5-215* root was consistently lower than that in the wild type background at different developmental stages (Supplemental Figure 11B-11D). In separately sampled shoots and roots, miR775 level determined by RT-qPCR was higher and lower in *hy5-215* compared to the wild type, respectively (Supplemental Figure 11E). These results indicate that *HY5* differentially regulates *MIR775A* in the aerial and underground organs.

**Figure 11.**
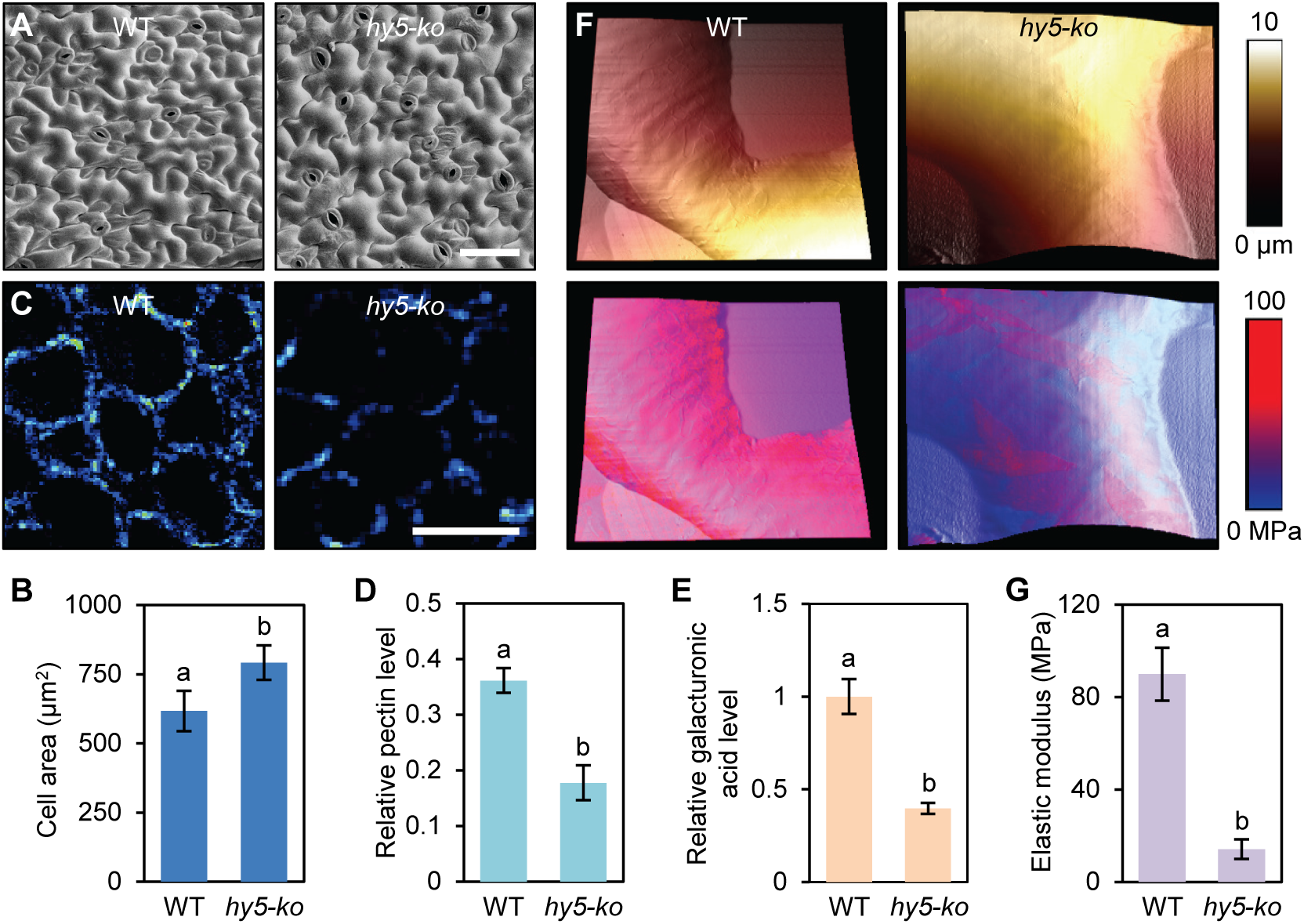
*HY5* Is a Negative Regulator of Leaf Size. (**A**) Enlargement of the *hy5-ko* epidermal cells in comparison to the wild type. The upper side of the fifth leaf from three-week-old plants was used for cryo-SEM analysis. Bar, 50 μm. (**B**) Quantification of epidermal cell size. Data are mean ± SD of 100 individual cells from five rosette leaves. Different letters denote significant difference (Student’s *t*-test, *p* < 0.001). (**C**) Imaging pectin in mesophyll cells by confocal Raman microscopy. Bar, 50 μm. (**D**) Average intensity of Raman images was used to deduce relative pectin levels. Data are mean ± SD of 15 areas from five leaves. Different letters denote significant difference (Student’s *t*-test, *p* < 0.01). (**E**) Quantification of the relative galacturonic acid level in the wild type and *hy5-ko* cell walls. Data are mean ± SD from three technical replicates performed on pooled leaves. Different letters denote significant difference (Student’s *t*-test, *p* < 0.01). (**F**) Topography of the wild type and *hy5-ko* cotyledon epidermal cells mapped by AFM (top) and cell topography overlaid with elasticity (bottom). (**G**) Quantification of apparent Young’s modulus. Each measurement was the average in a 5 μm by 5 μm area of a cell with the highest modulus. Data are mean ± SD of 10 cells from three cotyledons. Different letters denote significant difference (Student’s *t*-test, *p* < 0.001).

### The *HY5-MIR775A-GALT9* Pathway Regulates Leaf Size

The above findings prompted us to examine the role of *HY5* in leaf size determination. We generated a null *hy5-ko* allele by deleting almost the entire coding region using the CRISPR/Cas9 system (Supplemental Figure 12). Similar to the well-characterized *hy5-215* allele, which carries a point mutation that abolishes proper splicing of the first intron (Oyama et al., 1997), the *hy5-ko* seedlings exhibited larger cotyledons and longer hypocotyls than the wild type (Supplemental Figure 12B-12D). In the adult stage, the *hy5* mutants have larger rosette leaves and longer petioles than the wild type (Supplemental 12E). On the contrary, *HY5-OX* plants exhibited the opposite phenotypes in both the seedling and adult stages (Supplemental Figure 12C-12E). These results extended previous works documenting the organ enlargement phenotypes of the *hy5* mutants (Sibout et al., 2006; Brown and Jenkins, 2008; Burko et al., 2020).

**Figure 12.**
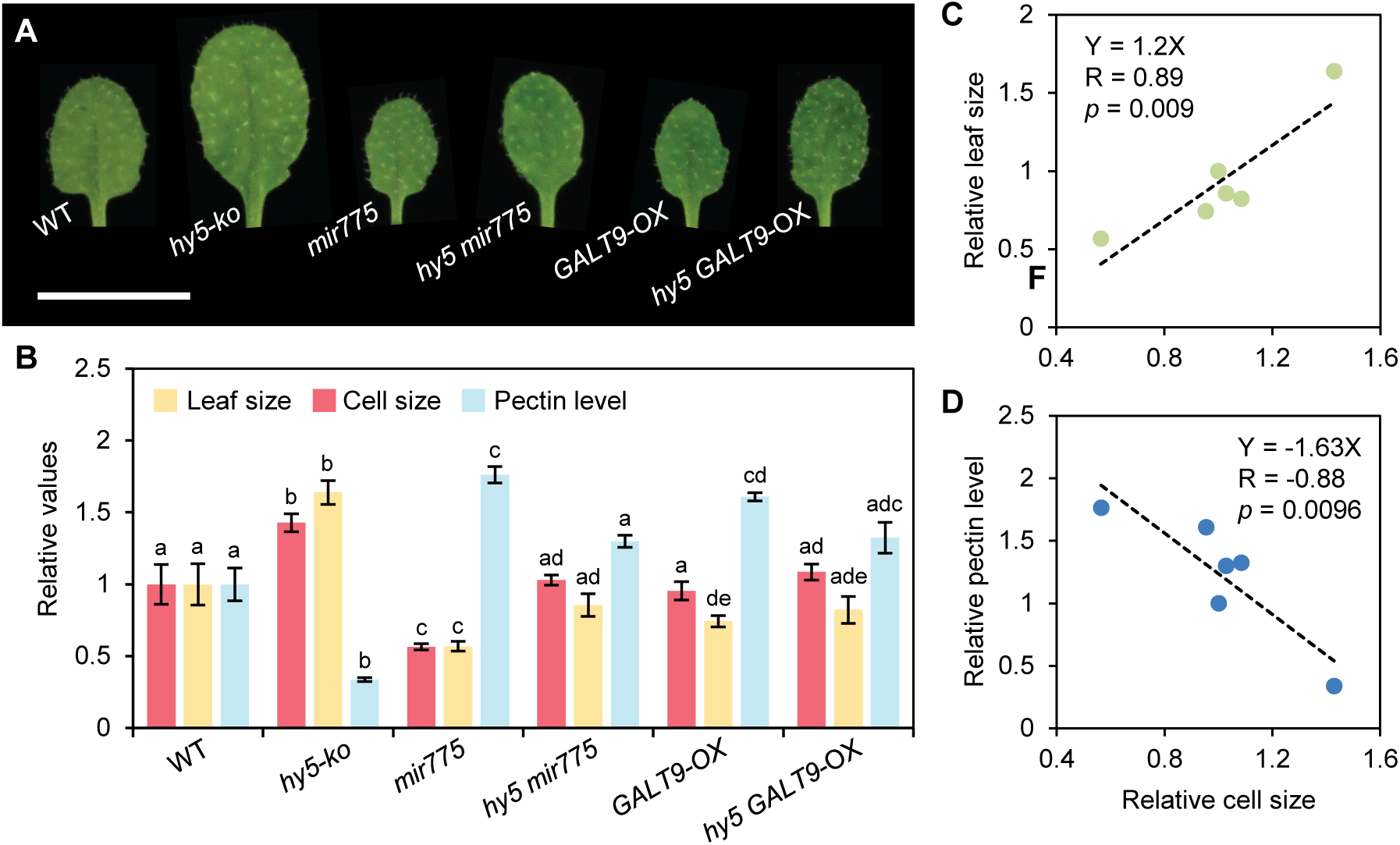
The *HY5-MIR775A-GALT9* Pathway Regulates Leaf Size. (**A**) Morphology of the fifth rosette leaves of three-week-old plants from the indicated genotypes. Bar, 5 mm. (**B**) Quantification of the leaf size, epidermal cell size, and pectin level relative to the wild type. Data are mean ± SD from 10 individual plants for leaf size, from 100 individual cells of several plants for cell size, and from three technical replicates performed on pooled leaves for galacturonic acid level. Within each set of measurements, different letters denote genotypes with significant difference (Student’s *t*-test, *p* < 0.05 for leaf size; *p* < 0.01 for cell size and galacturonic acid level). (**C-D**) Linear regression between cell sizes and organ sizes (C) and between cell sizes and galacturonic acid levels (D) across the six genotypes.

Using cryo-SEM, we analyzed and quantitated size of epidermal cells from both the cotyledons (Supplemental Figure 12F and 12G) and the fifth rosette leaves of adult plants (Figure 11). In both cases, we confirmed that the *hy5* mutants have significantly enlarged epidermal cells compared to the wild type. To test whether these effects were related to the pectin level, we performed Raman microscopy on the fifth rosette leaves and found that the *hy5-ko* cells have significantly less pectin than the wild type (Figure 11C and 11D). This finding was corroborated by quantifying the galacturonic acid content in the cell wall of the *hy5-ko* and wild type leaves (Figure 11E). AFM analysis showed that the *hy5-ko* cell walls have significantly reduced elastic modulus than the wild type (Figure 11F and 11G). These results indicate that *HY5* is a negative regulator for leaf size by increasing the pectin level and limiting cell expansion.

To genetically analyze whether *HY5* and *MIR775A-GALT9* act in the same pathway to regulate leaf growth, we generated the *hy5 mir775* and *hy5 GALT9-OX* double mutants through genetic crossing using *hy5-ko*. Quantification of the size of the fifth rosette leaves revealed that the leaf enlargement phenotype of *hy5-ko* was suppressed in both *hy5 mir775* and *hy5 GALT9-OX* (Figure 12). By cryo-SEM analysis and chemical quantification, we confirmed that the two double mutants mitigated the cell enlargement and pectin reduction phenotypes of *hy5-ko* (Figure 12B). Moreover, a linear correlation between the cell size and leaf size was observed for the wild type, *hy5-ko*, *mir775*, *GALT9-OX*, *hy5 mir775* and *hy5 GALT9-OX* genotypes (Figure 12C). Conversely, a reverse correlation between cell size and pectin level was observed across the same genotypes (Figure 12D). Taken together, these results indicate that *MIR775A* and *GALT9* act downstream of *HY5* in the same genetic pathway to control pectin content and intrinsic leaf size (Figure 13).

**Figure 13.**
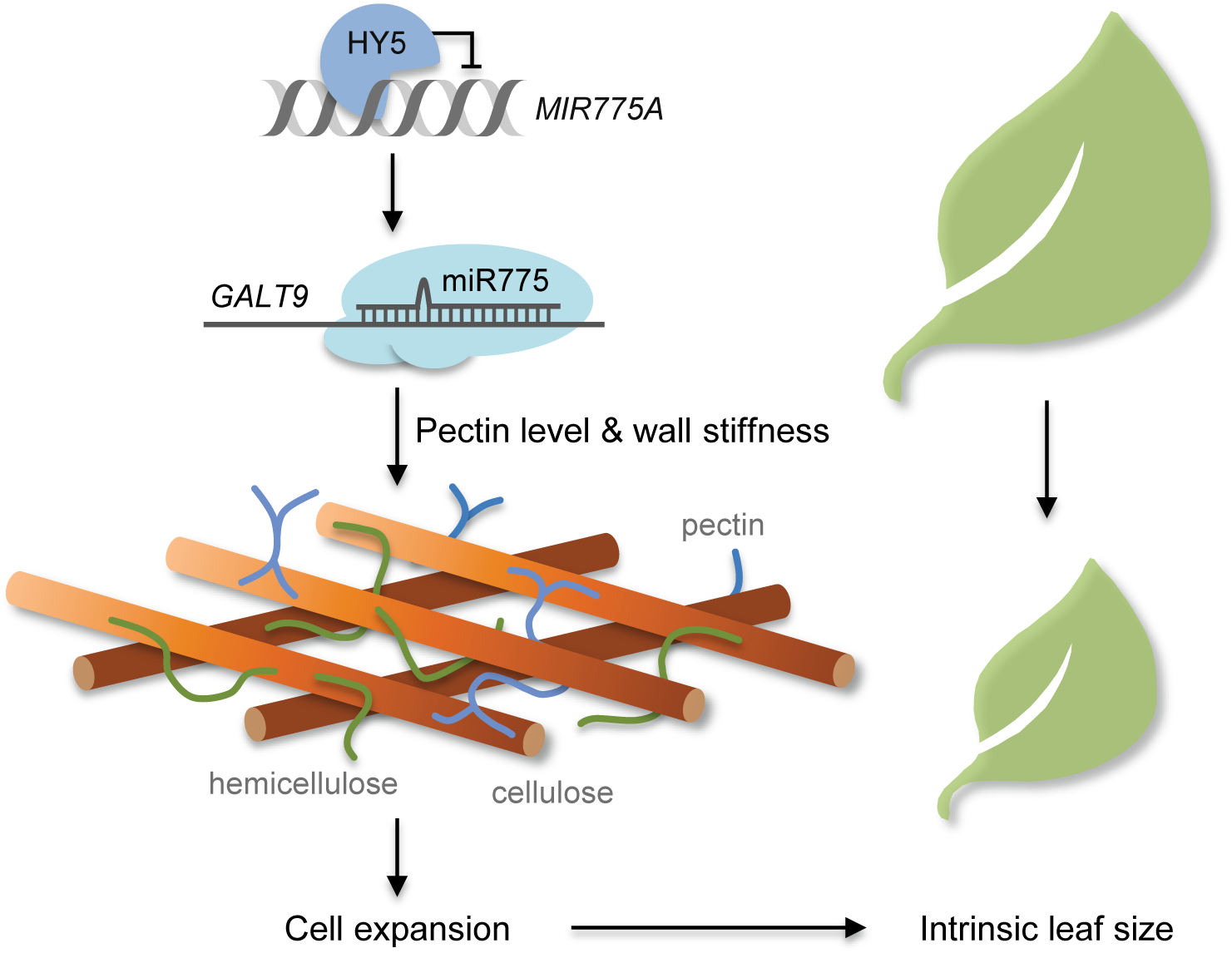
Model for the *HY5-MIR775A-GALT9* Pathway in Controlling Intrinsic Leaf Size. *HY5-MIR775A-GALT9* is a delineated double repression cascade for regulating GALT9 accumulation for leaf size determination. GALT9 participates in cell wall remodeling by promoting the pectin constituent and reducing cell wall elasticity, which may prepare the cells with proper resistance to turgor pressure for reaching the intrinsic size during leaf development.

## Discussion

Organ size is one of the dominating traits for plant development and architecture. Molecular genetics studies in the past three decades have identified numerous genes in organ size control (Bogre et al., 2008; Johnson and Lenhard, 2011; Gonzalez et al., 2012; Hepworth and Lenhard, 2014; Hong et al., 2018). Characterization of these genes has led to the conclusion that organ size control is primarily exerted by cell number regulation and cell size control is also integral to the intricate regulatory network governing organ size (Ferjani et al., 2007; Hong et al., 2018). Because the presence of a rigid plant cell wall, increasing of cell volume must be accompanied by mechanisms that allow timely wall relaxation. In this study, we identified 23 putative CW-miRNAs in *A. thaliana* that are potentially pertinent to the regulation of primary wall biosynthesis (Figure 1A). We selected miR775 as an example for functional characterization and provided new insights into how miRNAs may regulate organ size by modulating cell wall biosynthesis and/or modification.

We found that *GALT9* is the *bona fide* target for miR775 specifically in *A. thaliana* (Figures 1-3; Supplemental Figures 1 and 2). GALT9 is a member of the glycosyltransferase 31 family (Supplemental Figure 2A) and locates to the endomembrane (Figure 6A). It has been shown that several members of this family are capable of adding galactose to various glycans (Velasquez et al., 2011; Qin et al., 2017). The closest homolog to *GALT9* in cotton is *GhGALT1* (Supplemental Figure 2A). It was reported that *GhGALT1* overexpression in cotton resulted in smaller leaves, reduced boll size, and shorter fibers (Qin et al., 2017). *In vitro* purified GhGALT1 exhibited galactosyltransferase enzyme activity in galactan backbone biosynthesis (Qin et al., 2017). In this study, we provided a coherent body of evidence, including co-expression pattern with pectin related genes (Figure 6B-6D), monosaccharide quantification (Figure 7), confocal Raman microcopy and pectin immunolabelling (Figure 8; Supplemental Figure 10), that support an indisputable role of GALT9 in modulating the level of cell wall pectin in *A. thaliana*.

Moreover, reduction in pectin content in *galt9* is associated with alteration to cell wall mechanical property. Using AFM, we analyzed both the cotyledon and petal epidermal cells and observed that the *galt9* and *MIR775A-OX* cell walls displayed significantly lower elastic modulus than that of the wild type (Figure 9; Supplemental Figure 10). This observation is consistent with previous AFM analysis of epidermal cells that linked variation in the pectin network to changes in cell wall elasticity (Peaucelle et al., 2015; Xi et al., 2015). Together with studies on pectin biochemistry (Wolf et al., 2012; Xiao et al., 2014; Peaucelle et al., 2015; Andres-Robin et al., 2018), these findings suggest that attenuation of the pectin constitute in *galt9* and *MIR775A-OX* cell walls might compromise cross-link with cellulose, which in turn reduces elastic resistance to internal turgor pressure. This property of the cell wall would allow more expandability that translates into enlarged cell sizes, which we observed by cryo-SEM and AFM (Figures 5 and 9). Consistent with previous suggestions (e.g. Xiao et al., 2014), these results imply that the capacity for cell expansion is not maximized in the wild type organs due to rigidification of the pectin cross-linked cell walls. We hypothesize that by tuning pectin content, GALT9 might act as a downstream component of the regulatory networks that control cell expansion and present this idea in a conceptual model shown in Figure 13.

Regarding phyllome organs, we found that *MIR775A-OX* and *galt9* plants have significantly larger organs while *mir775* and *GALT9-OX* plants have smaller organs than the wild type (Figure 3; Supplemental Figures 3-7). Importantly, we did not observe substantial changes in the number of epidermal cells in any the examined organs (Figure 5). Across multiple organs of the *mir775*, *MIR775A-OX*, *galt9*, and *GALT9-OX* genotypes, a strong linear correlation between organ size and cell size was observed (Figure 5F). These changes in cell size resulted in essentially one-to-one changes in organ size across the examined genotypes (Figure 5F), suggesting that altered cell proliferation is not the cause for the observed changes in organ size. These findings thus indicate that the *MIR775A-GALT9* circuit is part of the cellular machinery that controls intrinsic organ size independent of cell proliferation (Ferjani et al., 2007; Hong et al., 2018).

Organogenesis requires coordinated cellular responses to developmental and environmental cues to realize the genetically determined growth potential. Through molecular and genetic analyses, we showed that in aerial organs *MIR775A* is under negative transcriptional control by *HY5* (Figure 10; Supplemental Figure 11). Extending previous studies (Sibout et al., 2006; Brown and Jenkins, 2008; Burko et al., 2020), we confirmed that *HY5* is a negative regulator for leaf size by modulating cell size (Figures 11 and 12; Supplemental Figure 12). Importantly, we found that the effect of *HY5* on cell size stems from alteration of pectin level and elasticity of the cell walls (Figures 11 and 12). *HY5-MIR775A-GALT9* is therefore a repression cascade operating in *A. thaliana* that imposes restriction on cell wall flexibility via GALT9-mediated pectin deposition and helps the plant to reach the desired intrinsic leaf size (Figure 13). HY5 is a key gene regulator for light signaling and photomorphogenesis (Oyama et al., 1997; Burko et al., 2020). Thus, whether the *HY5-MIR775A-GALT9* pathway is a mechanism for modulating pectin in the establishment of photomorphogenesis warrants investigation.

As *HY5* is a negative regulator of *MIR775A* (Figure 10), there should exist positive regulators for the spatiotemporal accumulation of miR775. Our preliminary results suggest that members of the class II *TCP* (*TEOSINTE BRANCHED1, CYCLOIDEA, PCF*) transcription factor family, which regulate the transition from cell division to cell expansion in dicot leaves (Palatnik et al., 2003; Ori et al., 2007; Efroni et al., 2008; Schommer et al., 2014), are candidates that activate *MIR775A*. It would be interesting to characterize these organogenesis-related factors that regulate miR775 to further elucidate how this miRNA contributes to pectin dynamics during leaf development. These efforts should be instrumental to reveal how other CW-miRNAs relay developmental or environmental cues to regulate cell wall remodeling and prepare the cells transitioning into expansion-driven growth with proper resistance to turgor pressure to reach the intrinsic size.

As an important class of endogenous regulatory RNAs, miRNAs are known to regulate leaf organogenesis (Palatnik et al., 2003; Ori et al., 2007; Rodriguez et al., 2010; Schommer et al., 2014; Rodriguez et al., 2016; Yang et al., 2018). Several conserved miRNA families, including miR156, miR319, and miR396, have been shown to regulate diverse aspects of leaf organogenesis involving leaf initiation, phase transition, polarity establishment, and morphology (Braybrook and Kuhlemeier, 2010; Efroni et al., 2010; Yang et al., 2018). For instance, over activation of miR319 promotes cell proliferation and results in larger leaves made up of smaller cells in comparison to the wild type (Palatnik et al., 2003; Efroni et al., 2008). These phenotypes are in line with the “compensation phenomenon” whereby mutants defective in cell proliferation may alter cell size to reach relatively the same final organ size (Ferjani et al., 2007; Kawade et al., 2010; Czesnick and Lenhard, 2015). Our finding on the role of miR775 in regulating leaf size through cell wall remodeling adds one more node to the miRNA networks governing leaf development and morphogenesis in *A. thaliana*.

The miRNA families with known roles in leaf organogenesis, such as miR156, miR319, and miR396, are deeply conserved in angiosperm (Yang et al., 2018; Guo et al., 2020). In contrast, while the target gene *GALT9* is conserved in angiosperm (Figure 1D; Supplemental Figure 2A), miR775 is an evolutionarily young miRNA unique to *A. thaliana* (Figure 1A; Supplemental Figure 1). Delineation of the *HY5-MIR775A-GALT9* pathway and documentation of the *mir775* phenotype (Figures 3-5, 10, and 12) demonstrated that *MIR775A* has been successfully integrated into the *A. thaliana* leaf developmental program. This finding suggests that the miRNA networks governing leaf development in different plant species may contain conserved “old” miRNAs interlaced with diverse species-specific “young” miRNAs. To confirm miRNA diversity in contributing to differential organ size control mechanisms, it would be interesting to test whether introducing species-specific CW-miRNAs such as miR775 or custom-designed artificial miRNAs into diverse plant species is sufficient to repress the *GALT9* orthologs and to modify organ size.

In summary, the evidence presented in this work highlights the function of a species-specific CW-miRNA in regulating cell and organ size in *A. thaliana*. Future investigation of other CW-miRNAs should provide additional insights into how plants orchestrate a complex sequence of molecular behaviors to modify the cell walls during development and in response to environmental cues. In addition to further elucidating the regulatory programs, these efforts would serve as a proof-of-concept to employ CW-miRNAs to sculpture plant size and architecture, which determine many agronomic traits in crops (Tang and Chu, 2017).

## Methods

### Plant Materials and Growth Conditions

The wild type plant used in this study was *A. thaliana* ecotype Col-0. To produce the *35S:MIR775A* and *35S:GALT9* constructs, the genomic regions containing pre-miR775a and the *GALT9* coding region were PCR amplified using the Pfusion DNA polymerase (New England Biolabs) and primers listed in Supplemental Table 2. The PCR products were cloned into the 35S-pKANNIBAL vector (Li et al., 2010). The *35S:GALT9^m^* construct was generated by substituting the nucleotides of the miR775 binding site within the *GALT9* coding region but not the encoded amino acids using primers listed in Supplemental Table 2. Following transformation and selection with BASTA (20 mg L^−1^) (bioWORLD), transformants were allowed to propagate to the T_2_ generation for analysis. The *HY5-OX* plants were as previously described (Gao et al., 2020). The *pMIR775A:GUS* line was generated by cloning the 1,064 bp genomic fragment upstream of the full-length cDNA BX81802 into the pCAMBIA-1381Xa vector (CAMBIA). The construct was used to transform wild type plants following the standard floral dipping method and selected with Hygromycin (20 mg L^−1^). T_2_ generation plants were screened for GUS activity and a designated line was used for crossing into the *hy5-215* background.

A CRISPR/Cas9 system specific for plants was used to delete *MIR775A*, *GALT9*, and *HY5* as described (Mao et al., 2013). In the modified pCAMBIA1300 vector, the *35S* and the *AtU6-26* promoter respectively drive *Cas9* and pairs of sgRNA designed to target both ends of the target genes. The resulting constructs were introduced into wild type plants via transformation. T_1_ generation plants were individually genotyped by PCR and sequencing to identify deletion events. Approximately 200 individual T_2_ generation plants were further genotyped to identify Cas9-free homozygous mutant lines.

To grow *Arabidopsis* plants, surface sterilized seeds were plated on agar-solidified MS media including 1% (w/v) sucrose and incubated at 4°C for three days in the dark. Germinated seedlings were either allowed to grow on the plate for three weeks (16 h light/8 h dark at 22°C/20°C) or transferred commercial soil and maintained in a growth chamber (16 h light/8 h dark at 22°C/20°C, 50% relative humidity). The light intensity was approximately 120 μmol m^−2^ s^−1^. Tobacco seedlings used for transient assay were *Nicotiana benthamiana*, which were grown under settings of 16 h light/8 h dark, 25°C/21°C, 50% relative humidity, and light intensity of 150-200 μmol m^−2^ s^−1^.

### Identification of CW-miRNAs

The 572 cell wall biosynthesis related genes were collected by GO term search. The 491 genes encoding Golgi-enriched proteins were obtained from previous studies (Parsons et al., 2012). Full-length cDNA sequences for a nonredundant combination of these genes were obtained from TAIR (www.arabidopsis.org). Searching against the 427 annotated miRNAs in *A. thaliana* (miRBase, version 22) (Kozomara et al., 2018) was done using the computational tools psRNATarget (Dai and Zhao, 2011) and psRobot (Wu et al., 2012). This process produced two separate outputs, which were further searched against degradome sequencing data generated by the CleaveLand4 or StarScan pipeline (Addo-Quaye et al., 2009; Liu et al., 2015). Possible miRNA-target pairs predicted by both tools or by either one but compatible with degradome data were combined into a nonredundant dataset, which contained 23 miRNAs and 78 target genes listed in Supplemental Table 1. Conservation of CW-miRNAs was determined by searching against miRNAs in miRBase (version 22) and PmiREN (Guo et al., 2020). *Brassicaceae* species with genome sequences but no miRNA annotation were manually checked using BLASTN (E-value < 1e^−10^) and RNAfold for evaluating the secondary structures as previously reported (Gruber et al., 2008). The predicted target genes were searched against seven Brassicaceae species with sequenced genomes for possible orthologs using BLASTP (E-value < 1e^−10^).

### Degradome Sequencing and Analysis

Total RNA from *MIR775A-OX* leaves was isolated using Trizol reagent (Invitrogen). Degradome library construction using biotinylated random primers was performed as previously described (German et al., 2008; 2009). The library was subjected to single-end sequencing (50 bp) on the Illumina Hiseq 2500 platform. A total of 63,558,618 clean reads were generated and 55,077,460 mapped to the TAIR10 *A. thaliana* genome using Bowtie2 (Langmead and Salzberg, 2012), allowing no more than two mismatches. The sequencing data were deposited to the Sequence Read Archive database (SRR10322040). Three sets degradome sequencing data from the wild type seedlings (SRR3945024, SRR3945025, and SRR3945026) were used as control. Reads mapped to the predicted target sites were used to extrapolate the positions of the 5’ transcript ends and to calculate the RPM values using an in-house Perl script.

### Quantitative RNA Analyses

Total RNA was isolated using the Quick RNA Isolation kit (Huayueyang). Each sample was taken from the pooled tissues, such as leaves or roots. All experiments were repeated on at least three sets of independently prepared RNA. mRNA and miRNA were reverse transcribed into cDNA using the SuperScript III reverse transcriptase (Invitrogen) and the miRcute Plus miRNA First-Stand cDNA Synthesis kit (Tiangen), respectively. Quantitative PCR was performed with the ABI PRISM Fast 7500 Real-Time PCR engine using the TB Green Premix Ex TaqII (TIi RNaseH Plus) (TaKaRa) and the miRcute Plus miRNA qPCR kit (SYBR Green) (Tiangen) with three technical replicates, respectively. *Actin7* and 5S RNA were used as internal controls. Relative amounts of mRNA and miRNA were calculated using the comparative threshold cycle method.

### 5’ RLM-RACE

The assay was performed using the 5’-Full RACE kit (TaKaRa) according to the manufacturer’s instructions with modifications. Total RNA was isolated from seedlings and ligated to the 5’ RNA adaptor by T4 RNA ligase (TaKaRa). Reverse transcription was performed with 9-nt random primers and the cDNA amplified by PCR with an adaptor primer and a gene-specific primer. This was followed by a nested PCR and cloning of the products using the Mighty TA-cloning kit (TaKaRa). Twenty independent clones were randomly picked and sequenced.

### REN/LUC Dual Luciferase Assays

The *REN/LUC* construct was modified from the previous version (Liu et al., 2014) by using the *Actin2* promoter to drive the LUC fusion proteins. The *GALT9^m^-LUC* reporter construct was generated by substituting the nucleotides in the miR775 binding site within *GALT9* by PCR using primers listed in Supplemental Table 2.Three combinations of the two effectors and/or reporter constructs were used to transiently co-transform tobacco protoplasts as previously described (Liu et al., 2014). Chemiluminescence was detected using the NightSHADE LB 985 system (Berthold) in the presence of 20 mg mL^−1^ potassium luciferin (Gold Biotech). The LUC/REN ratio was calculated to infer effectiveness of miR775 targeting.

### Protein Localization

The GALT9 and RAN1 coding sequences were respectively cloned into the pJIM19-GFP/mCherry/ vectors. *Agrobacterium* GV3101 cells harboring the *35S:GALT9-GFP* and *35S:RAN1-mCherry* constructs were mixed and co-infiltrated into tobacco leaf epidermal cells with a syringe. The cells were observed three days thereafter using an LSM 710 laser scanning confocal microscope (Zeiss). Colocalization was analyzed using the Coloc 2 module in ImageJ.

### Co-expression Analysis

The *GALT9* co-expressed genes in *A. thaliana* were obtained from the ATTED-II database (version 9) (Obayashi et al., 2018). The 174 co-expressed genes were identified based on the mutual rank index as a co-expression measure using a cutoff value of 400. The co-expressed genes were visualized using the built-in tools in ATTED-II.

### Cryo-SEM

The method for cryo-SEM was as previously described (Esch et al., 2004) with minor modifications. The scanning electron microscope FEI Helios NanoLab G3 UC (Thermo Scientific) and the Quorum PP3010T workstation (Quorum Technologies), which has a cryo preparation chamber connected directly to the microscope, were used as a unit. Plant samples were frozen in subcooled liquid nitrogen (−210°C) and then transferred in vacuum cabin to the cold stage of the chamber for sublimation (−90°C, 5 min) and sputter coating (10 mA, 30 sec) with platinum. Images were taken using the electron beam at 2 kV and 0.2 nA with a working distance of 4 mm. Projective cell area of indicated samples was measured using ImageJ. Average cell size was determined by measuring 100 cells from at least three samples.

### Chemical Analysis of Cell Wall Components

Cell wall cellulose level was determined using the Cellulose Extraction and Determination kit (Comin Biotechnology, www.cominbio.com). Approximately 300 mg tissues per sample were homogenized in 1 mL 80% ethanol, heated at 90°C for 20 min, cooled to room temperature, and centrifuged at 6000*g* for 10 min. The insoluble pellets were washed once in 1 mL 80% ethanol and once in 1 mL acetone by vertexing and centrifugation at 6000*g* for 10 min. The pellets were resuspended in 1 mL solution I provided in the kit, de-starched for 15 h at room temperature, and collected by centrifugation at 6000*g* for 10 min, and dried. Five milligrams of the resulting cell wall materials were homogenized in 0.5 mL distilled water, mixed with 0.75 mL concentrated sulfuric acid on ice, incubated for 30 min, and centrifuged at 8000*g* for 10 min at 4°C. Glucose determination in the supernatants was based on the anthrone assay (Yuan et al., 2019; Huang et al., 2020) using reagents provided in the kit and following the manufacturer’s protocol. The glucose concentration from the blue-green samples was measured by absorbance at 630 nm using a NanoPhotometer P-class USB spectrophotometer (Implen GmbH).

Pectin level was determined using the Pectin Extraction and Determination kit (Comin Biotechnology). Briefly, approximately 50 mg tissues per sample were homogenized in 1 mL extraction buffer I provided in the kit, heated at 90°C for 30 min, cooled to room temperature, and centrifuged at 5000*g* for 10 min. The insoluble pellets were washed in 1 mL extraction buffer I by vertexing and centrifugation at 5000*g* for 10 min. The pellets were resuspended in 1 mL extraction buffer II provided in the kit, heated at 90°C for 1 h, and centrifuged at 8000*g* for 15 min. Galacturonic acid in the supernatants was determined by colorimetry as previously described (Taylor, 1993) using reagents provided in the kit. Absorbance of the pink- to red-colored samples at 530 nm was read on the NanoPhotometer P-class USB spectrophotometer.

### GUS Staining

Care was taken to make sure whole plants or seedlings were submerged and evenly incubated at room temperature for 6 h in a GUS staining solution (1 mM 5-bromo-4-chloro-3-indolyl-b-D-glucuronic acid, 100 mM Na_3_PO4 buffer, 3 mM each K_3_Fe(CN)_6_/K_4_Fe(CN)_6_, 10 mM EDTA, and 0.1% Nonidet P-40). After staining, chlorophyll was removed using 70% ethanol for 4 h, which was repeated three times.

### Confocal Raman Imaging

Freshly detached *Arabidopsis* cotyledons and young leaves were washed sequentially with 70%, 100%, and 70% ethanol for 10 min each to remove chlorophyll. After that, the samples were kept in water. Label-free imaging of cellulose and pectin was performed with a home-built coherent Raman microscope, fitted with a picoEmerald (Applied Physics & Electronics) picosecond laser as light source, which supplies tunable pump beam and fixed Stokes beam. As previously described (Gierlinger et al., 2012), 1100 cm^−1^ (asymmetric stretching vibration of the glycoside bond C-O-C) and 854 cm^−1^ (C-O-C skeletal mode of α-anomers) were used for specific *in situ* mapping of cellulose and pectin, respectively. The pump beams were respectively tuned to 952.5 nm and 975.5 nm, synchronized, and visualized with an inverted microscope (Olympus) equipped with a 25× objective lens and a coherent Raman detection module. Each image was acquired with 512 by 512 pixels and averaged by 5 frames. A background image was acquired for each sample by only illuminating with the pump laser beam. For normalization, difference of the signal intensity between each image and the corresponding background image was divided by the background image using ImageJ.

### Pectin Immunolabelling

This procedure was performed as previously described (Qi et al., 2017). Briefly, seven-day-old seedlings were fixed in absolute methanol under vacuum and embedded in Steedman’s wax (Sigma-Aldrich). After rehydration, 8 μm sections were prepared and pre-treated for 1 h with 2% (w/v) BSA in PBS, and then incubated overnight with the primary antibody LM19 (PlantProbes) diluted 1:500 in 0.1% BSA. After three washes in BST buffer (0.1% BSA and 0.1% (v/v) Tween 20), sections were incubated for 1 h with the secondary antibody Alexa Fluor 546 goat anti-rat IgG (Life Technologies) diluted 1:1,000 in 0.1% BSA. Sections were mounted in ProLong Antifade (Life Technologies) with cover slips and the Fluorescent Brightener 28 dye solution (Sigma-Aldrich) added. Fluorescence imaging was performed with an LSM 710 laser scanning confocal microscope (Zeiss).

### AFM Analysis

Freshly detached cotyledons and petals were subject to AFM analysis as described with modifications (Peaucelle et al., 2015; Xi et al., 2015). Briefly, the samples were attached to glass slide using transparent nail polish and submerged under water at room temperature to prevent plasmolysis. The topographical images of epidermal cells were scanned with a BioScope Resolve atomic force microscope equipped with a ScanAsyst-Fluid cantilever (Bruker) of 20 nm tip radius and 0.7 N m^−1^ spring constant. For topography, peak force error and DMT modulus images, Peak Force QNM mode of the acquisition software were used, with peak force frequency at 2 kHz and peak force set-point at 3 nN. The topology image size was 10 × 10 μm^2^ or 20 × 20 μm^2^ with a resolution of 256 × 256 pixels recorded at a scan rate of 0.2 Hz. To map apparent Young’s modulus, 1 to 2 mm-deep indentations were performed along the topological skeletons of epidermal cells to ensure relative normal contact between the probe and sample surface. At least three indentation positions were chosen for each cell, with each position consecutively indented three times, making at least nine indentation force curves per cell. Data were analyzed with Nanoscope Analysis version 1.8.

## Supplemental Data

Supplemental Figure 1. Comparison of Pre-miR775a Homologs in *A. thaliana* and *A. lyrata*.

Supplemental Figure 2. MiR775 Specifically Targets *GALT9* in *A. thaliana*.

Supplemental Figure 3. Characterization of *MIR775A-OX* Lines.

Supplemental Figure 4. Generation and Characterization of the *mir775* Mutant Lines.

Supplemental Figure 5. Characterization of the *MIR775A-OX mir775* Line.

Supplemental Figure 6. Generation and Characterization of the *galt9* Mutant Lines.

Supplemental Figure 7. Characterization of the *GALT9-OX* Lines.

Supplemental Figure 8. Degradome Sequencing Profiles of Predicted MiR775 Targets.

Supplemental Figure 9. Phenotypic Comparison of the *galt9* and *dcl1* Mutants.

Supplemental Figure 10. Analysis of the *qrt2* Mutant Defective in Pectin Turnover.

Supplemental Figure 11. H*Y*5 Differentially Regulates *MIR775A* in the Shoot and the Root.

Supplemental Figure 12. Generation and Characterization of Mutants for *HY5*.

Supplemental Table 1. Putative CW-miRNAs and Predicted Target Genes in *A. thaliana*.

Supplemental Table 2. Oligonucleotide Sequences of the Primers Used in This Study.

Supplemental Dataset 1. *GALT9* Co-expressed Genes in *A. thaliana*.

## Accession Number

Sequence data from this article can be found in the *Arabidopsis* Genome Initiative or GenBank/EMBL databases under the following accession numbers: *MIR775A* (At1g78206), *HY5* (At5g11206), *GALT9* (At1g53290), *DCL1* (At1g01040), and *QRT2* (At3g07970). T-DNA insertion mutants used are *galt9* (SALK_015338), *dcl1* (SALK_056243C), and *qurt2* (SALK_031337).

## Author Contributions

L.L. designed and supervised the research. H.Z., Y.Z., J.D., J.P., L.L, T.W., and H.C. performed the research. H.Z., Y.S., Z.G. (Guo), Z.G. (Gao), L.X., G.Q., and Y.J. analyzed the data. H.Z. and L.L. wrote the paper.

## Acknowledgements

We thank Drs. Dong Liu and Chan Li at the National Center for Protein Science at Peking University for technical assistance in AFM operation and image analysis, Dr. Yiqun Liu and Ms. Yifeng Jiang at the Core Facilities of School of Life Sciences at Peking University for assistance with SEM. This work was supported by grants from the National Key Research and Development Program of China (2017YFA0503800) and the National Natural Science Foundation of China (31621001).

---

**Supplemental Table 1.** Putative CW-miRNAs and Predicted Target Genes in *A. thaliana*.

| **MiRNA** | **Target** | **Description** |
| --- | --- | --- |
| miR156h | AT5G38610 | PECTIN METHYLESTERASE INHIBITOR |
| miR156i | AT1G13560 | AMINOALCOHOLPHOSPHOTRANSFERASE1 |
|  | AT3G01390 | VACUOLAR MEMBRANE ATPASE10 |
|  | AT5G38610 | PECTIN METHYLESTERASE INHIBITOR SUPERFAMILY PROTEIN |
| miR156j | AT2G33040 | GAMMA SUBUNIT OF MITOCHONDRIAL ATP SYNTHASE |
|  | AT5G38610 | PECTIN METHYLESTERASE INHIBITOR SUPERFAMILY PROTEIN |
| miR1886 | AT1G02800 | GLYCOSIDE HYDROLASE FAMILY9 |
|  | AT2G36870 | XYLOGLUCAN ENDOTRANSGLYCOSYLASE/HYDROLASE |
| miR2936 | AT1G15690 | INORGANIC H PYROPHOSPHATASE FAMILY PROTEIN |
|  | AT1G15690 | PYROPHOSPHATE-ENERGIZED INORGANIC PYROPHOSPHATASE |
| miR414 | AT1G09210 | CALRETICULIN 1B |
|  | AT1G56340 | CALRETICULIN 1A |
|  | AT2G16600 | ROTAMASE CYP3 |
|  | AT3G25520 | RIBOSOMAL PROTEIN L5 |
|  | AT4G33740 | MYB-LIKE PROTEIN X |
|  | AT5G12110 | ELONGATION FACTOR 1-BETA 1 |
|  | AT5G13850 | NASCENT POLYPEPTIDE-ASSOCIATED COMPLEX SUBUNIT ALPHA-LIKE PROTEIN3 |
|  | AT5G61790 | CALNEXIN1 |
|  | AT4G33330 | GLUCURONYLTRANSFERASE |
|  | AT2G31210 | BHLH TRANSCRIPTION FACTOR |
|  | AT3G50240 | KINESIN-RELATED PROTEIN |
| miR417 | AT5G66460 | ENDO-BETA-MANNANASE |
| miR4227 | AT4G12650 | ENDOMEMBRANE PROTEIN 70 FAMILY |
| miR4239 | AT3G57330 | AUTOINHIBITED Ca<sup>2+</sup>-ATPASE11 |
| miR5015 | AT1G71040 | LOW PHOSPHATE ROOT2 |
| miR5021 | AT1G09330 | ECHIDNA GOLGI APPARATUS MEMBRANE PROTEIN-LIKE PROTEIN |
|  | AT1G10950 | TRANSMEMBRANE NINE1 |
|  | AT1G11310 | SEVEN TRANSMEMBRANE MLO FAMILY PROTEIN |
|  | AT1G11680 | CYTOCHROME P450 51G1 |
|  | AT1G71940 | SNARE ASSOCIATED GOLGI PROTEIN FAMILY |
|  | AT2G18840 | INTEGRAL MEMBRANE YIP1 FAMILY PROTEIN |
|  | AT2G20120 | CONTINUOUS VASCULAR RING |
|  | AT2G26680 | FKBM FAMILY METHYLTRANSFERASE |
|  | AT3G08550 | ELONGATION DEFECTIVE1 |
|  | AT3G09440 | HEAT SHOCK PROTEIN 70 FAMILY PROTEIN |
|  | AT3G21160 | ALPHA-MANNOSIDASE2 |
|  | AT3G26370 | O-FUCOSYLTRANSFERASE FAMILY PROTEIN |
|  | AT3G49310 | MAJOR FACILITATOR SUPERFAMILY PROTEIN |
|  | AT3G52300 | ATP SYNTHASE D CHAIN |
|  | AT4G30190 | H(+)-ATPASE2 |
|  | AT4G30440 | UDP-D-GLUCURONATE 4-EPIMERASE1 |
|  | AT4G34180 | CYCLASE FAMILY PROTEIN |
|  | AT5G20350 | TIP GROWTH DEFECTIVE1 |
|  | AT5G51570 | SPFH/BAND 7/PHB DOMAIN-CONTAINING MEMBRANE-ASSOCIATED PROTEIN |
|  | AT5G23870 | PECTIN ACETYLESTERASE FAMILY PROTEIN |
|  | AT5G26670 | PECTIN ACETYLESTERASE FAMILY PROTEIN |
|  | AT3G26370 | O-FUCOSYLTRANSFERASE FAMILY PROTEIN |
|  | AT1G24170 | GALACTURONOSYLTRANSFERASE |
|  | AT4G36160 | NAC-DOMAIN TRANSCRIPTION FACTOR |
|  | AT5G33290 | XYLOGALACTURONAN XYLOSYLTRANSFERASE |
|  | AT4G02130 | GALACTURONOSYLTRANSFERASE |
|  | AT5G61130 | CALLOSE BINDING |
|  | AT1G53000 | NUCLEOTIDE-DIPHOSPHO-SUGAR TRANSFERASES SUPERFAMILY PROTEIN |
| miR5628 | AT2G02860 | SUCROSE TRANSPORTER2 |
| miR5630b | AT1G33120 | RIBOSOMAL PROTEIN L6 FAMILY |
| miR5658 | AT1G14670 | ENDOMEMBRANE PROTEIN 70 FAMILY |
|  | AT1G32090 | EARLY-RESPONSIVE TO DEHYDRATION4 |
|  | AT3G27220 | GALACTOSE OXIDASE/KELCH REPEAT SUPERFAMILY PROTEIN |
|  | AT3G47670 | PLANT INVERTASE/PECTIN METHYLESTERASE INHIBITOR SUPERFAMILY PROTEIN |
|  | AT4G11220 | VIRB2-INTERACTING PROTEIN2 |
|  | AT5G55500 | BETA-1,2-XYLOSYLTRANSFERASE |
|  | AT5G57655 | XYLOSE ISOMERASE FAMILY PROTEIN |
|  | AT1G05310 | PECTIN LYASE-LIKE SUPERFAMILY PROTEIN |
|  | AT2G06850 | ENDOXYLOGLUCAN TRANSFERASE EXGT-A1 |
|  | AT3G54920 | PECTATE LYASE-LIKE PROTEIN |
|  | AT4G29230 | NAC DOMAIN CONTAINING PROTEIN75 |
|  | AT1G62760 | PECTIN METHYLESTERASE INHIBITOR |
|  | AT1G20190 | ALPHA-EXPANSIN FAMILY PROTEIN |
|  | AT3G06260 | GALACTURONOSYLTRANSFERASE |
|  | AT5G62380 | NAC-DOMAIN TRANSCRIPTION FACTOR |
|  | AT3G62660 | GALACTURONOSYLTRANSFERASE |
| miR5662 | AT3G49010 | BREAST BASIC CONSERVED1 |
| miR773b | AT2G26890 | GRAVITROPISM DEFECTIVE2 |
| miR775 | AT1G53290 | GALACTOSYLTRANSFERASE |
| miR776 | AT2G32530 | CELLULOSE SYNTHASE |
| miR827 | AT1G63010 | VACUOLAR PHOSPHATE TRANSPORTER1 |
| miR837 | AT5G24810 | ABC1 FAMILY PROTEIN |
| miR838 | AT1G43170 | RIBOSOMAL PROTEIN1 |
|  | AT1G51630 | O-FUCOSYLTRANSFERASE FAMILY PROTEIN |
|  | AT1G51630 | PRENYLATED RAB ACCEPTOR 1.B1 |
| miR854e | AT3G56110 | PRENYLATED RAB ACCEPTOR1 |
| miR861 | AT3G58730 | VACUOLAR ATP SYNTHASE SUBUNIT D |
|  | AT1G71990 | LEWIS-TYPE ALPHA 1,4-FUCOSYLTRANSFERASE |

**Supplemental Table 2.**
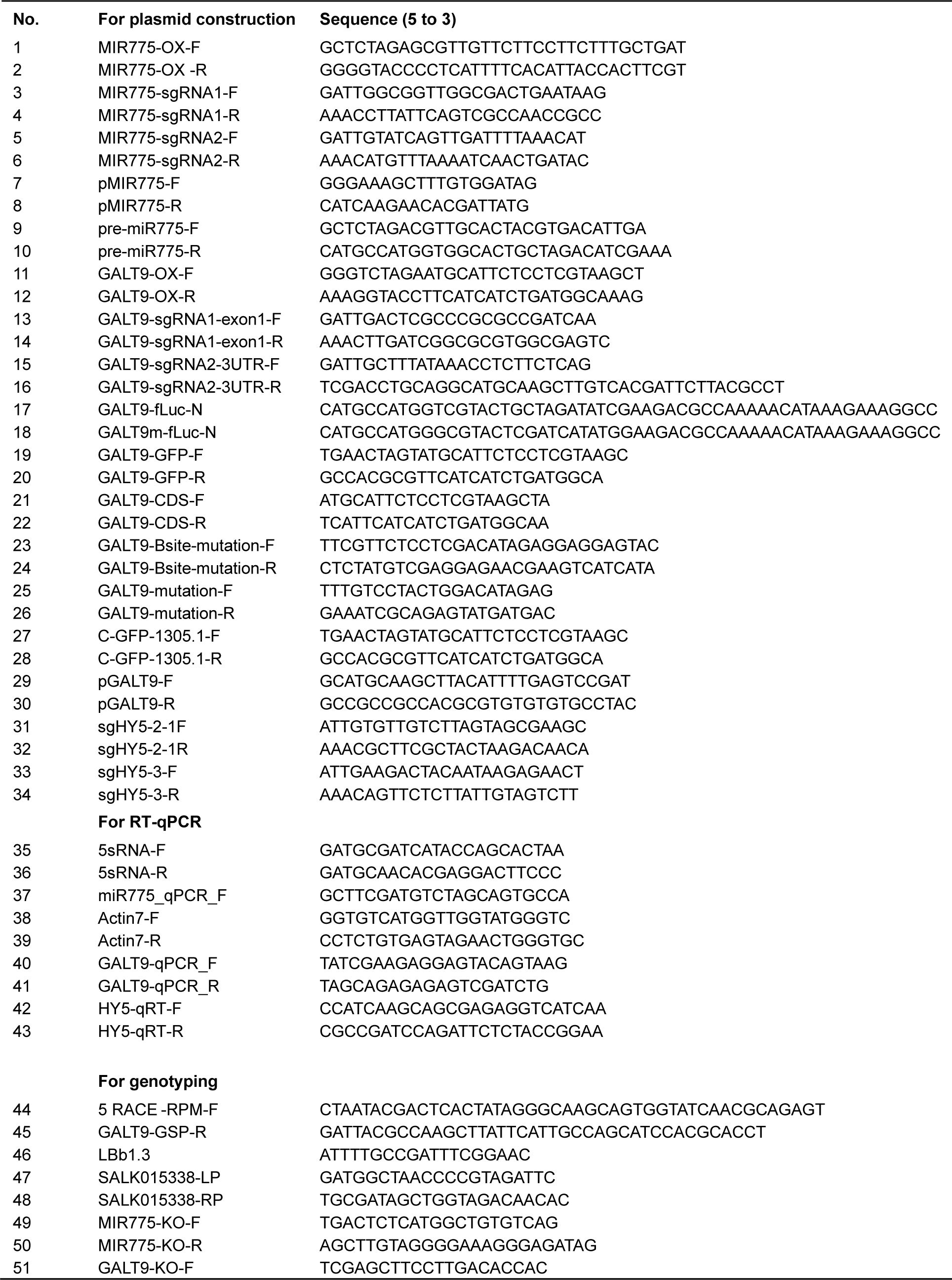
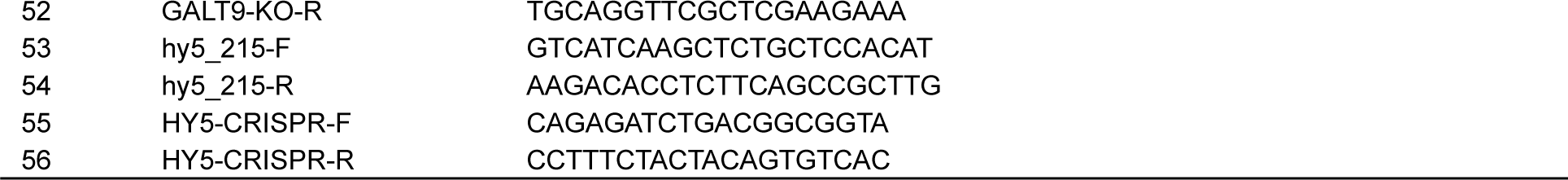
Oligonucleotide Sequences of the Primers Used in This Study.

**Supplemental Figure 1.**
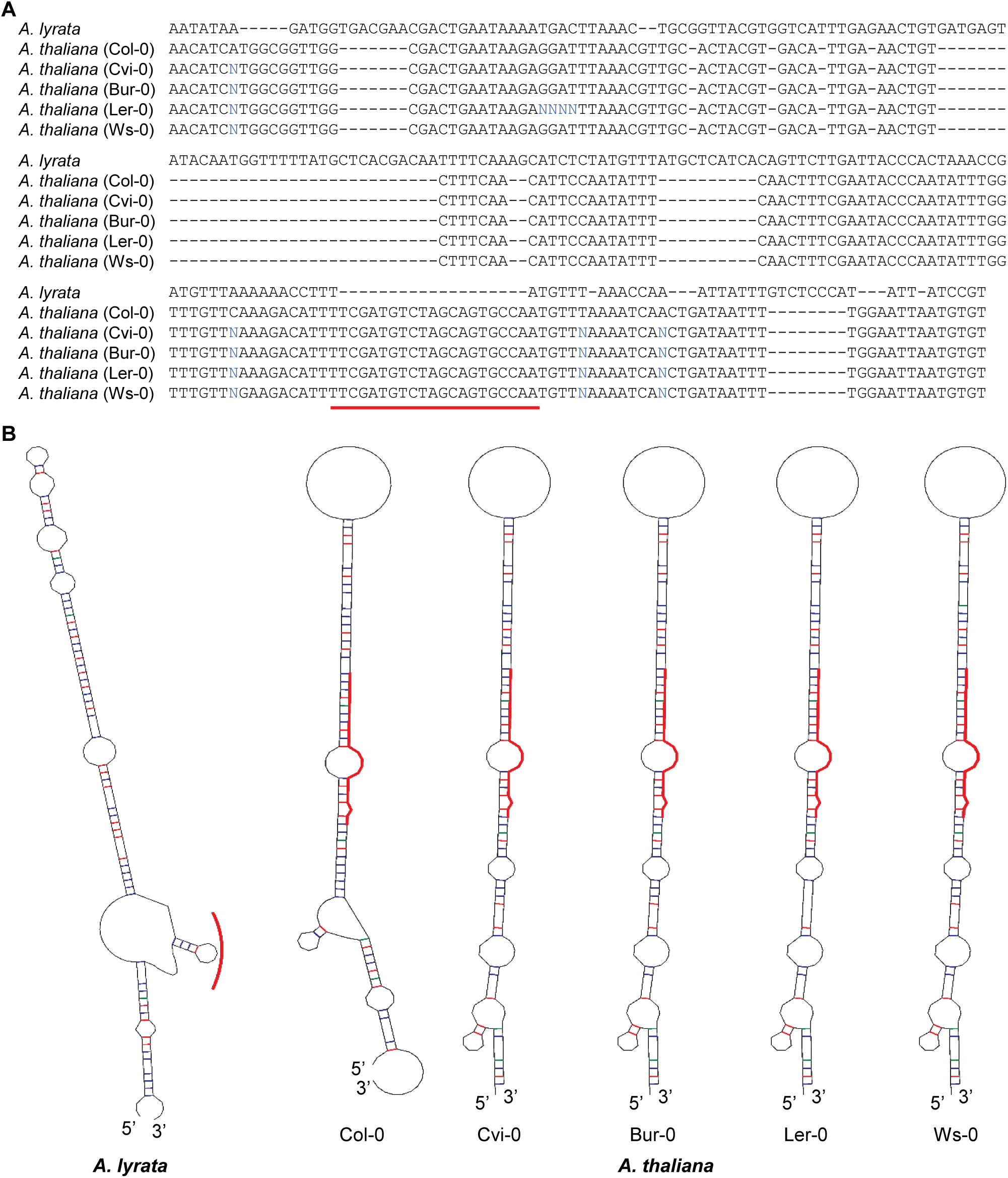
Comparison of Pre-miR775a Homologs in *A. thaliana* and *A. lyrata*. (**A**) Alignment of pre-miR775a sequences from five representative *A. thaliana* ecotypes with the closest homolog in *A. lyrata*. Sequences are 29,422,419-29,422,603 on *A. thaliana* (Col-0) chromosome 1 and 18,060,424-18,060,639 on *A. lyrate* chromosome 2. Region corresponding to mature miR775 is underlined in red. (**B**) Predicted secondary structures from sequences in A. Red lines indicate the region corresponding to miR775 in *A. thaliana*. Supports Figure 1 in the main manuscript.

**Supplemental Figure 2.**
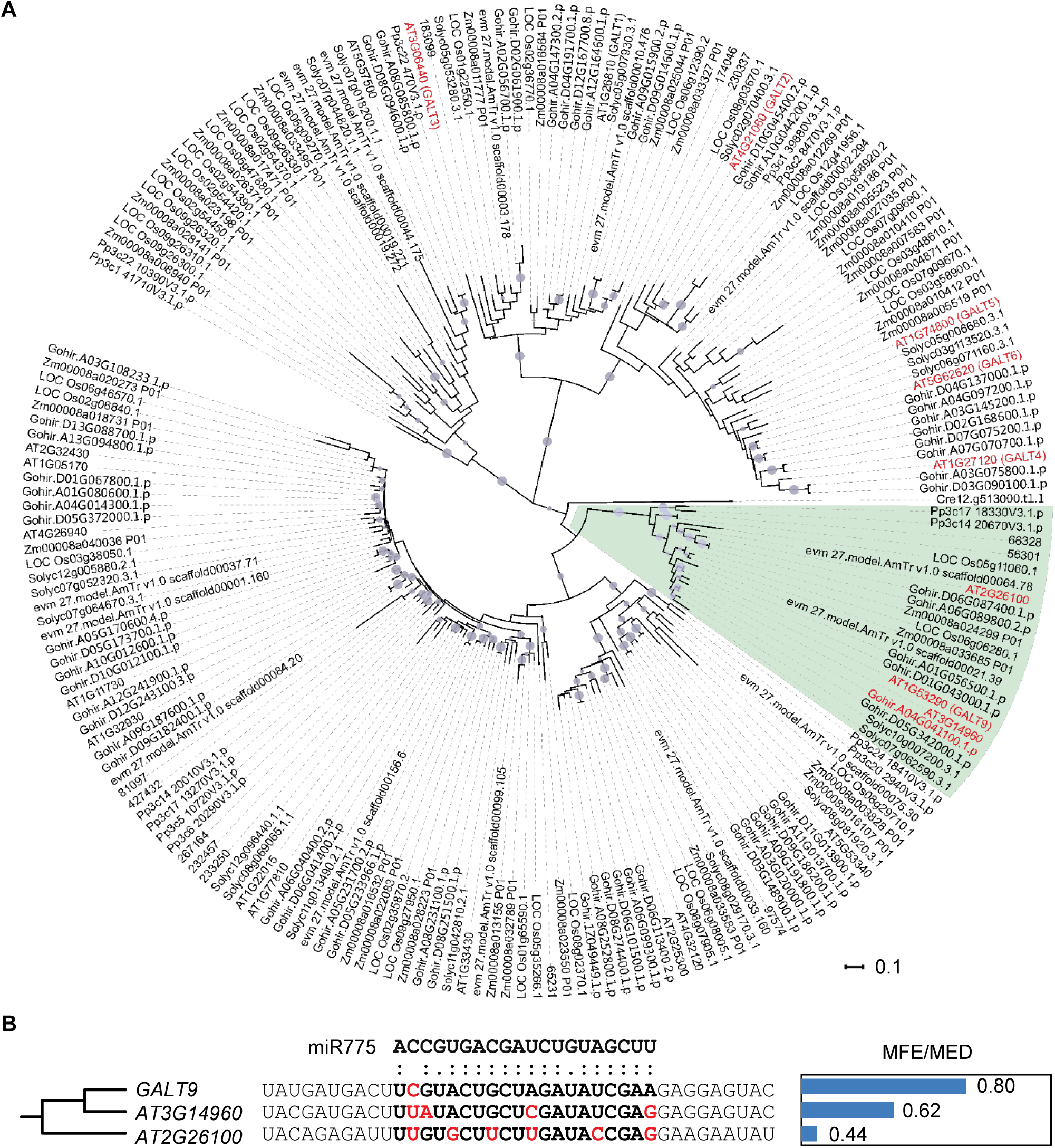
MiR775 Specifically Targets *GALT9* in *A. thaliana*. (**A**) Phylogeny of representative members of the glycosyltransferase 31 family. Shown is an unrooted neighbor joining tree built with the JTT model. Bootstrap values are from 1,000 iterations. Circles indicate branches with a bootstrap value > 60. The clade containing *GATL9* is shade in green. Genes known for involvement in primary cell wall biosynthesis are highlighted in red. (**B**) Sequence alignment at the miR775 binding site, shown in bold, between *GALT9* and two closest homologs in *A. thaliana*. Nucleotides undermining complementarity with miR775 are shown in red. The MFE/MED ratios are shown on the right, which indicate that only *GALT9* is a potential target for miR775. Supports Figure 1 in the main manuscript.

**Supplemental Figure 3.**
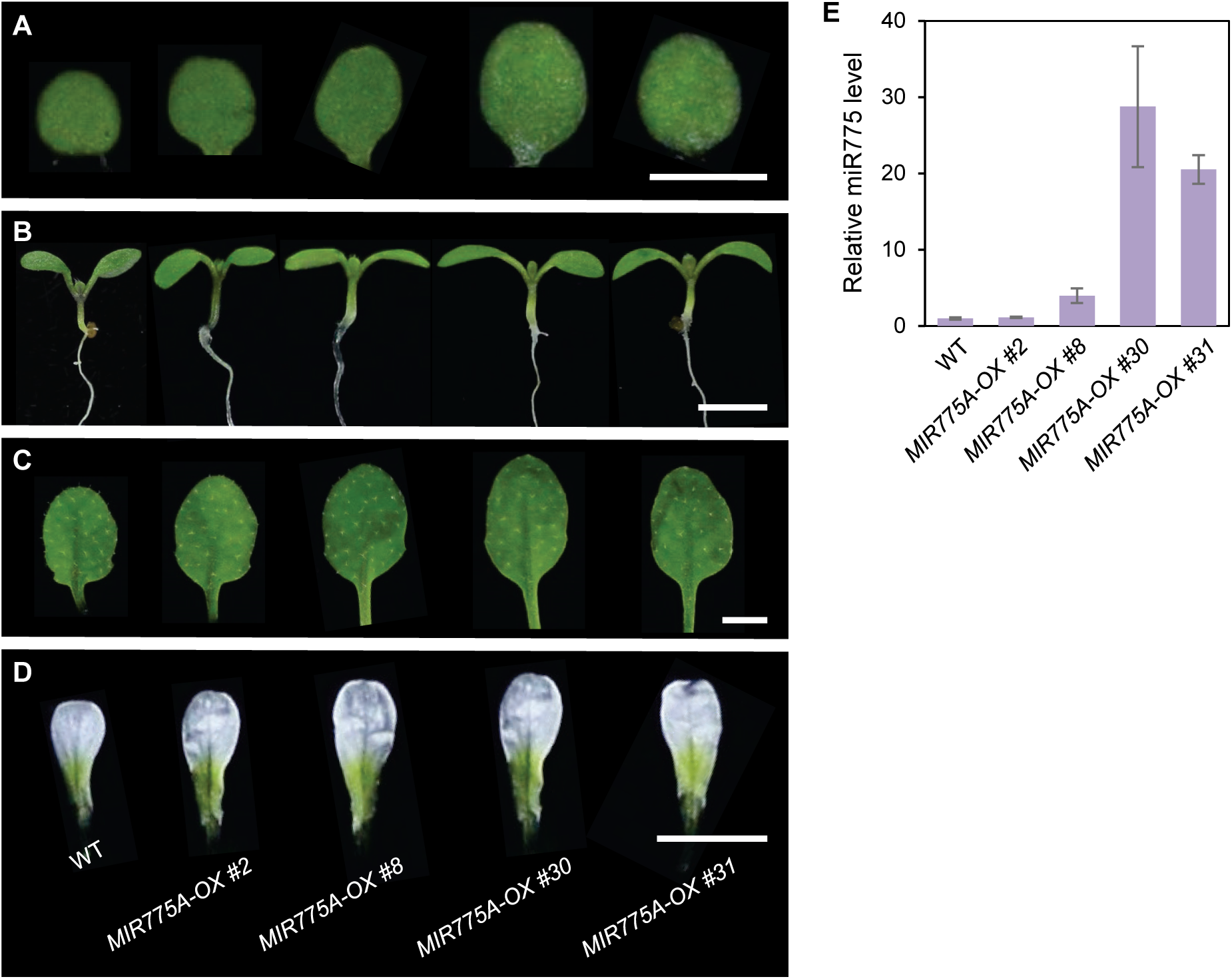
Characterization of *MIR775A-OX* Lines. (**A-D**) Morphological comparison of the indicated lines. Shown are cotyledon of eight-day-old seedlings (A), seedling showcasing the hypocotyl (B), the fifth rosette leaf of three-week-old plants (C), and petal of open flowers (D). Bars, 2 mm. *MIR775A-OX* was generated by expressing the *35S:pre-miR775a* transgene (pre-miR775a under control of the enhanced *35S* promoter) in *A. thaliana*. Seventeen independent T_1_ lines were obtained and four further analyzed at the T_2_ generation. Line #8 was selected for subsequent analyses. (**E**) RT-qPCR analysis of relative miR775 abundance in the selected lines. Data are means ± SD from three technical replicates. Supports Figures 2-4 in the main manuscript.

**Supplemental Figure 4.**
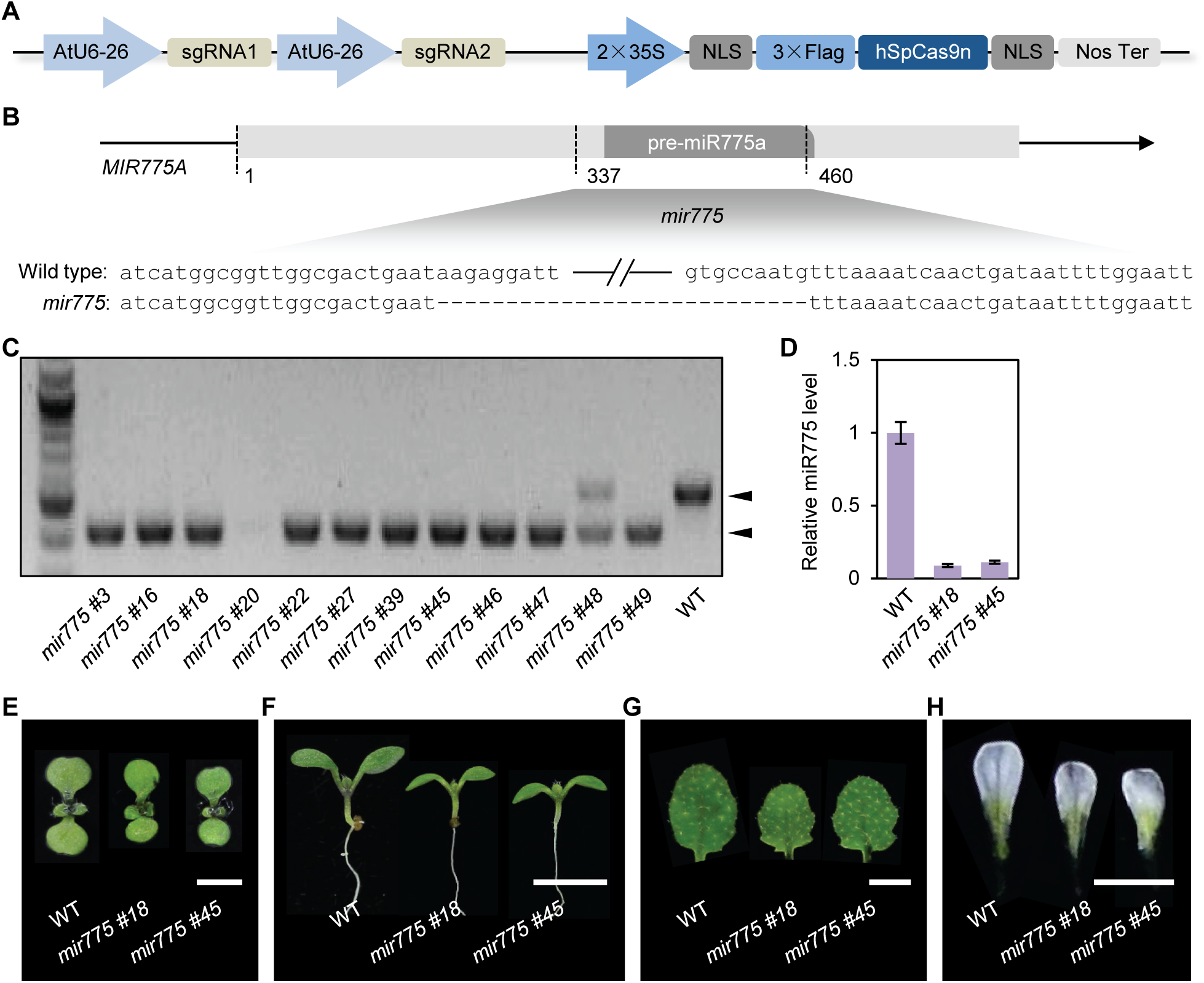
Generation and Characterization of the *mir775* Mutant Lines. (**A**) Diagram showing the CRISPR/Cas9 vector for simultaneously introducing Cas9 with paired sgRNAs. (**B**) Scheme for generating *mir775* deletion using the CRISPR/Cas9 system. Numbers mark positions according to the full length cDNA *BX818024*. The paired sgRNAs are designed to delete a 123 bp region encompassing pre-miR775a. Sequence comparison for a typical deletion allele with reference to the wild type allele is shown on the bottom. (**C**) Genotyping result for 10 independent homozygous *mir775* lines. Genomic DNA from individual deletion lines was PCR-amplified and gel-separated. Size polymorphisms according to the wild type and deletion alleles are indicated. Lines #18 and #45 were selected for subsequent analyses. (**D**) RT-qPCR analysis of relative miR775 abundance in the two selected lines. Data are means ± SD from three technical replicates. (**E-H**) Morphological comparison of the indicated lines. From left to right: eight-day-old seedlings showcasing the cotyledon (E), seedlings showcasing the hypocotyl (F), the fifth rosette leaves of three-week-old plants (G), and petals of open flowers (H). Bars, 2 mm. Supports Figures 3 and 4 in the main manuscript.

**Supplemental Figure 5.**
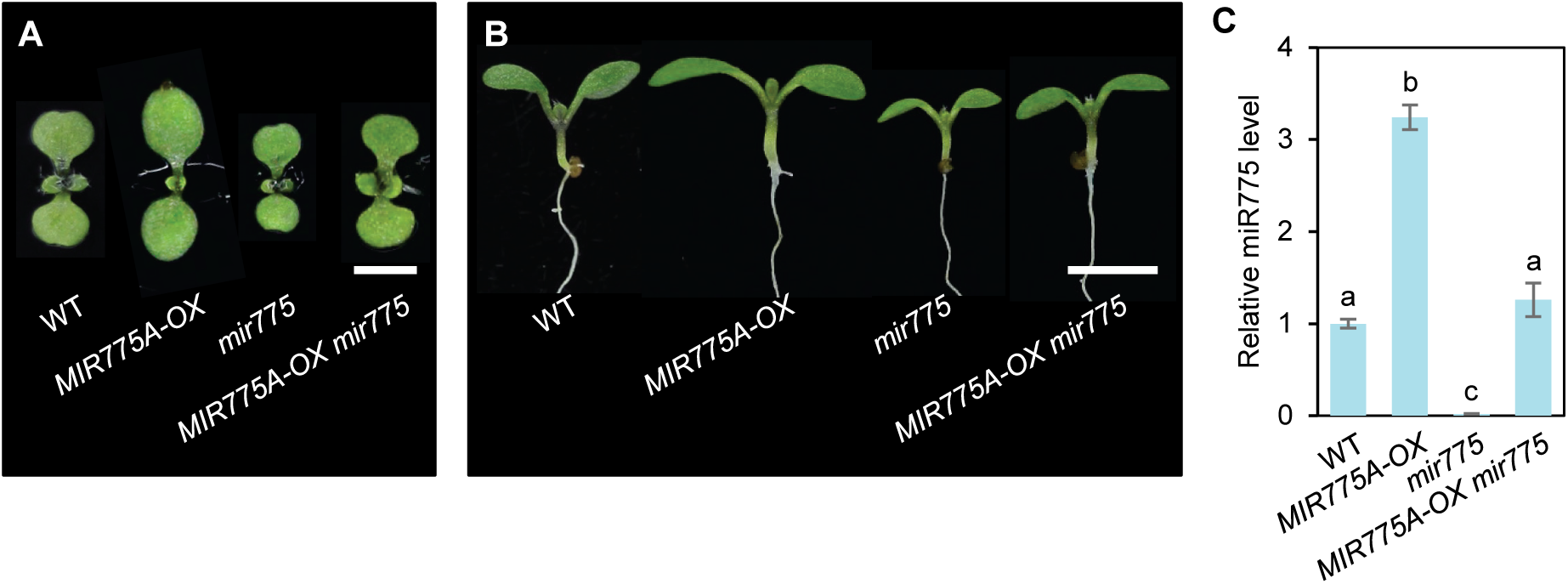
Characterization of the *MIR775A-OX mir775* Line. (**A-B**) Morphological comparison of the indicated genotypes. Eight-day-old seedlings were photographed to showcase the cotyledon (A) and the hypocotyl (B). *MIR775A-OX mir775* was created by crossing T_3_ generation *MIR775A-OX* line #8 to *mir775.* F_2_ progenies homozygous for *mir775* and resistant to BASTA (*MIR775A-OX* positive) were selected for analyses. Bars, 2 mm. (**C**) RT-qPCR analysis of relative miR775 abundance in the indicated genotypes. Data are means ± SD from three technical replicates. Different letters denote groups with significant difference (Student’s *t*-test, *p* < 0.001). Supports Figures 3 and 4 in the main manuscript.

**Supplemental Figure 6.**
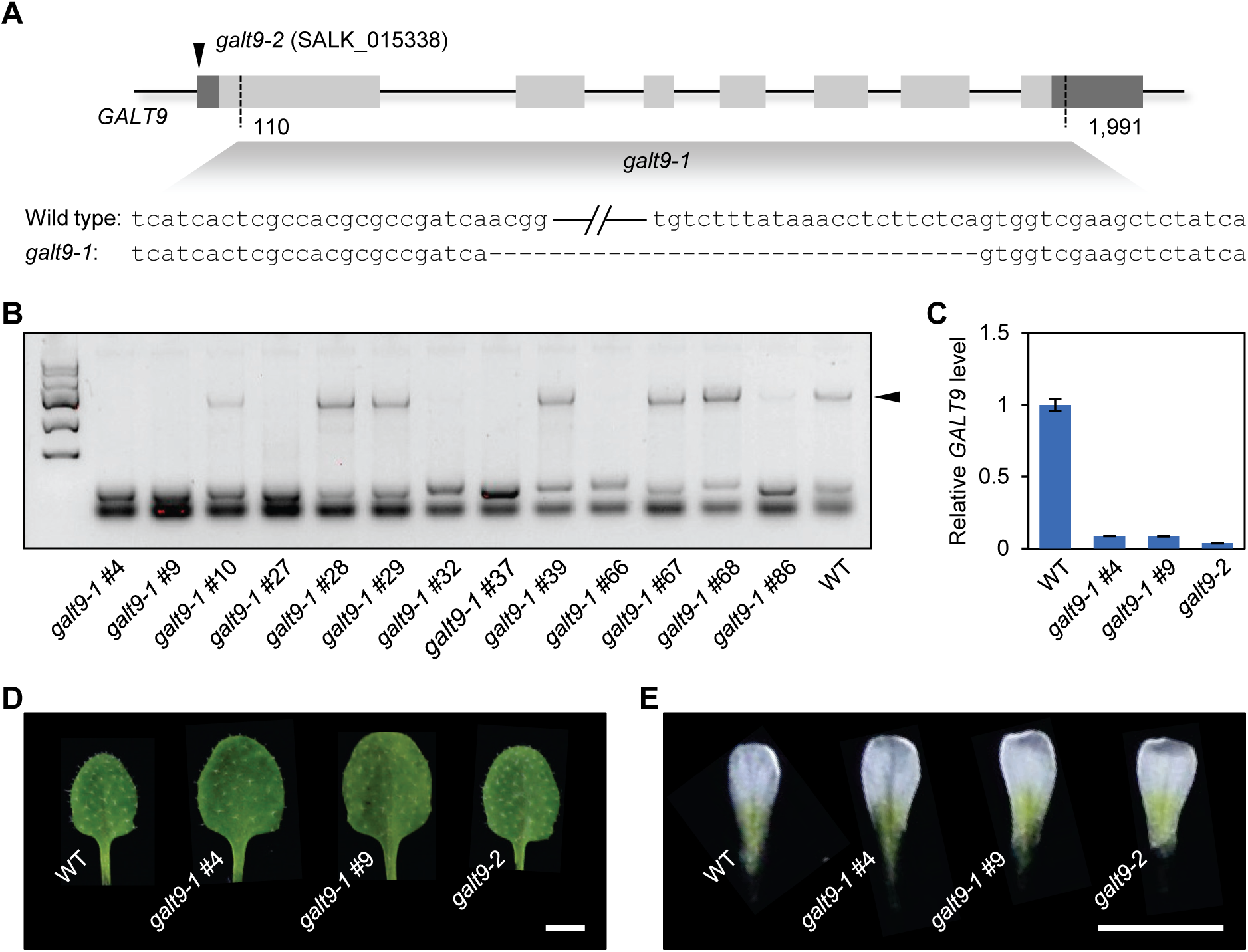
Generation and Characterization of the *galt9* Mutant Lines. (**A**) Scheme for generating *galt9* deletion mutants using the CRISPR/Cas9 system. Exons of *GALT9* are shown as horizontal boxes. Two sgRNAs are designed to create paired cleavage sites positioned at 110 and 1,991, resulting in a 1,882 bp deletion. The corresponding mutant was named *galt9-1*. A T-DNA insertion line (SALK_015338) with the T-DNA inserted into the start codon was named *galt9-2*. (**B**) Genotyping result for the deletion lines. A total of seven independent homozygous lines were identified. PCR product corresponding to the wild type allele is marked. Lines #4 and #9 were selected for subsequent analyses. (**C**) RT-qPCR analysis of relative *GALT9* transcript levels in the indicated lines in comparison to the wild type. Data are means ± SD from three technical replicates. (**D-E**) Morphology of the fifth rosette leaf (D) and petal (E) of the indicated genotypes. Bars, 2 mm. Supports Figures 3 and 4 in the main manuscript.

**Supplemental Figure 7.**
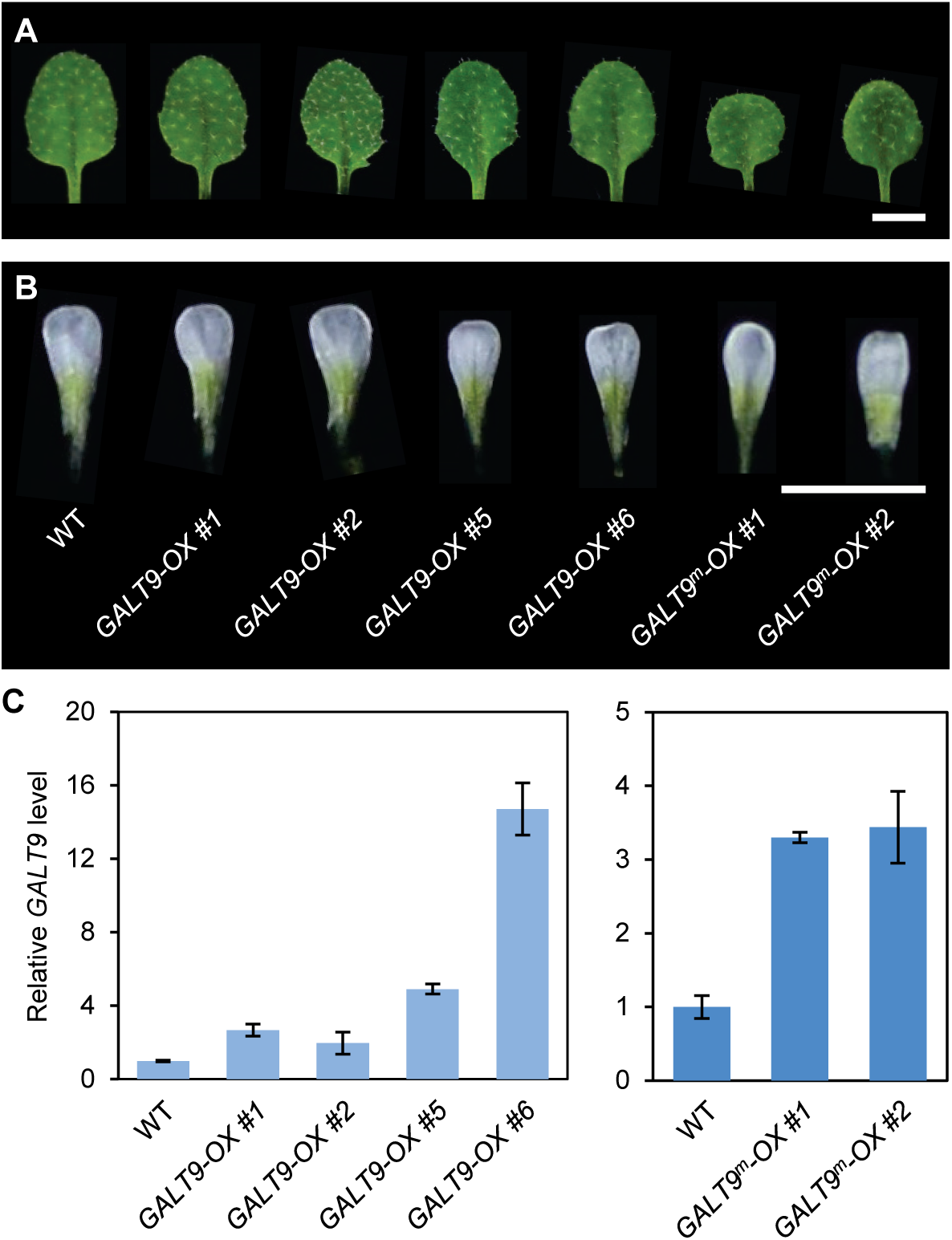
Characterization of the *GALT9-OX* Lines. (**A-B**) Morphological comparison of the fifth rosette leaf from three-week-old plants (A) and petal from open flowers (B). Bars, 2 mm. *GALT9-OX* was generated by expressing the *GALT9* coding region under control of the enhanced *35S* promoter in *A. thaliana*. Twelve independent T_1_ lines were obtained and four further analyzed at the T_2_ generation. *GALT9^m^-OX* was generated by substituting the nucleotides of the miR775 binding site in *GALT9* but not the encoded amino acids. Six independent T_1_ lines were obtained and two further analyzed at the T_2_ generation. (**C**) RT-qPCR analysis of relative *GALT9* transcript levels in the indicated lines in comparison to the wild type. Data are means ± SD from three technical replicates. Supports Figures 3 and 4 in the main manuscript.

**Supplemental Figure 8.**
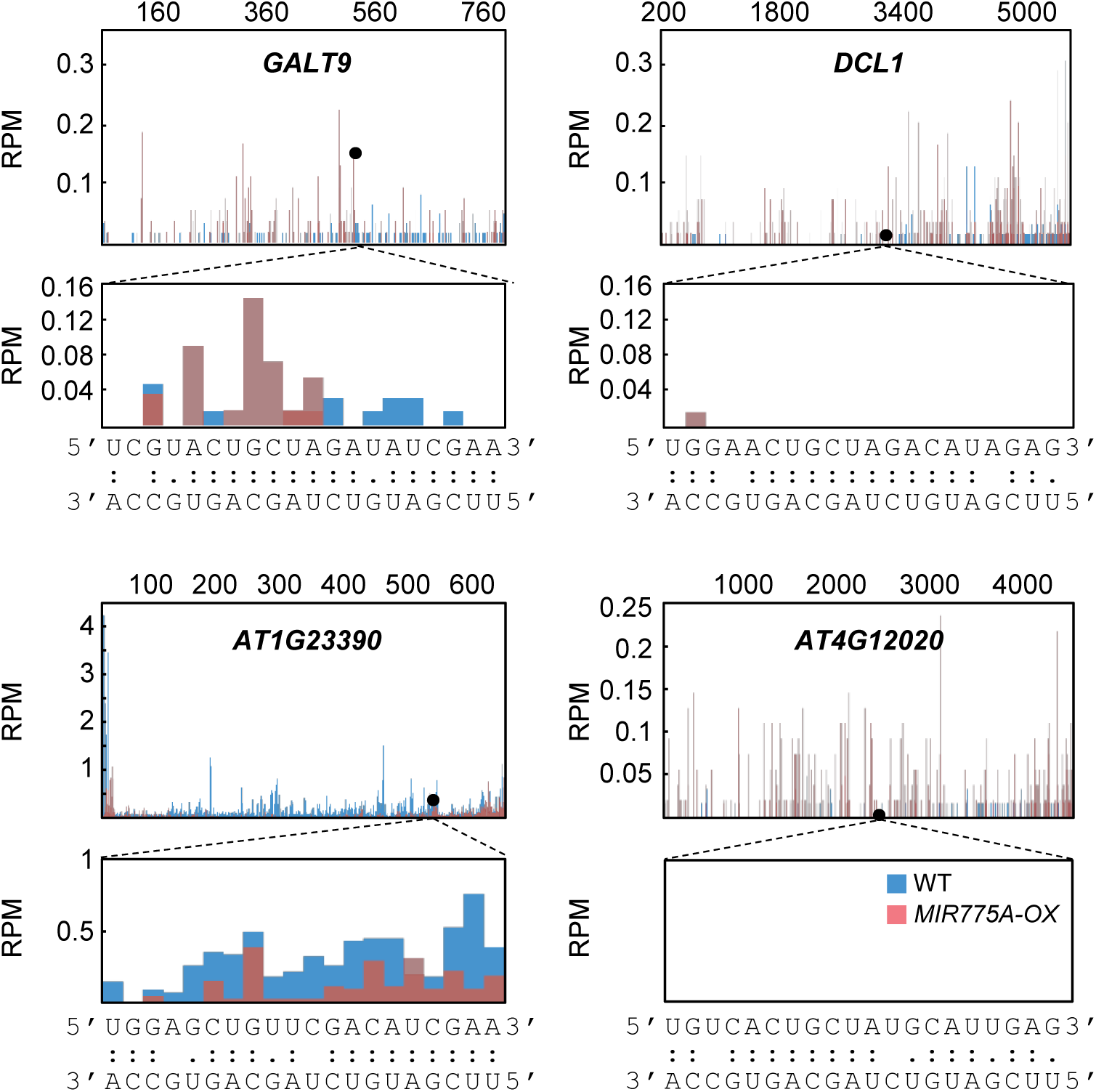
Degradome Sequencing Profiles of Predicted MiR775 Targets. Degradome sequencing data were obtained from the wild type and *MIR775A-OX* plants. Shown on top are normalized frequencies of reads with unique 5’ ends mapped to the four potential miR775 target genes. Enlarged views at the predicted miR775-binding sites are shown on the bottom along with base pairing pattern to miR775. Supports Figure 2 in the main manuscript.

**Supplemental Figure 9.**
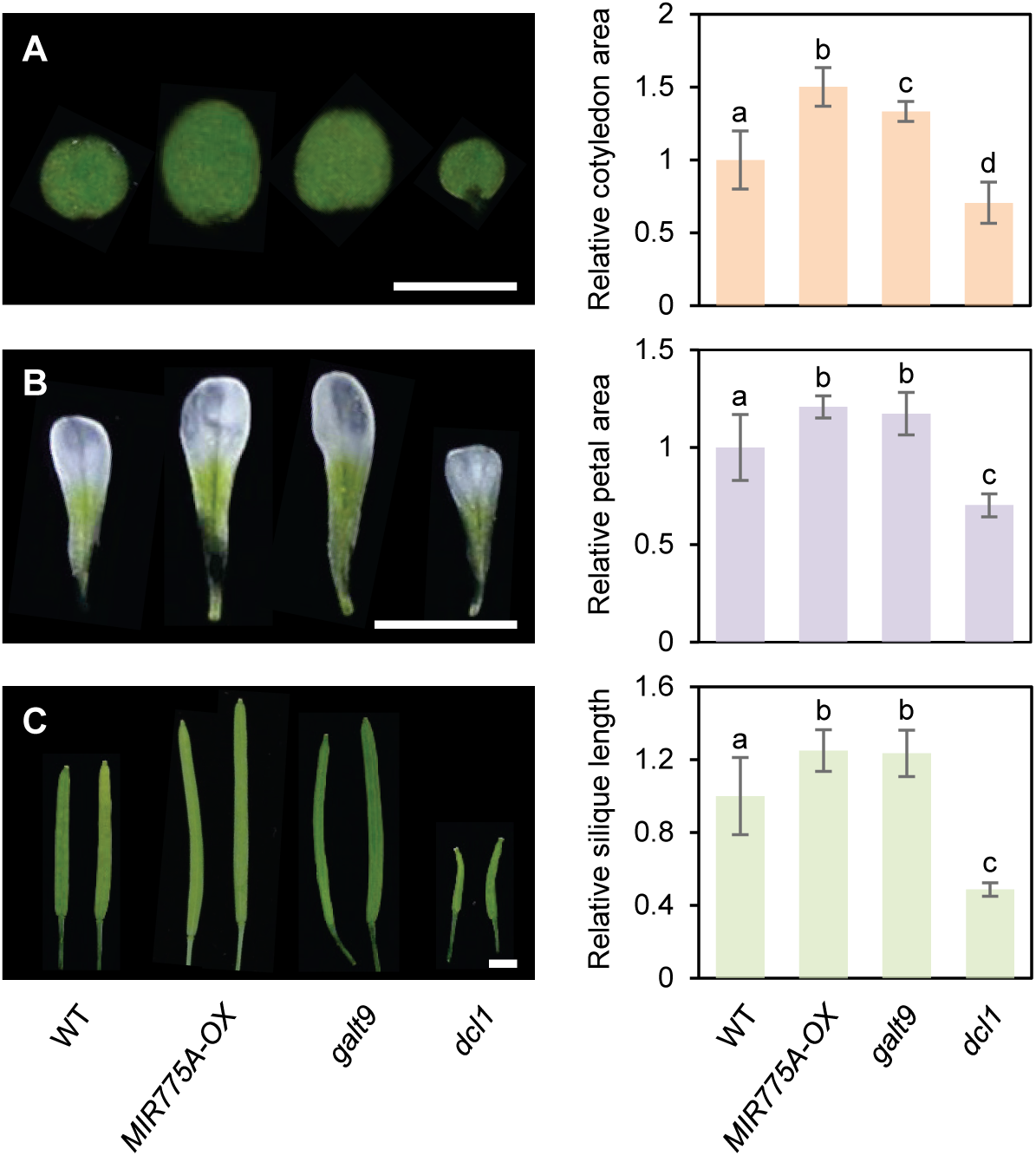
Phenotypic Comparison of the *galt9* and *dcl1* Mutants. (**A-C**) Morphological comparison of the indicated genotypes. Photographs of eight-day-old cotyledons (A), petals of open flowers (B), and mature siliques (C) are shown on the left. Bars, 2 mm. Quantifications of the relative cotyledon area, petal area, and silique length are shown on the right. Data are means ± SD from 30 individual organs normalized to the wild type. Different letters denote genotypes with significant difference (Student’s *t*-test, *p* < 0.01). Supports Figures 2-4 in the main manuscript.

**Supplemental Figure 10.**
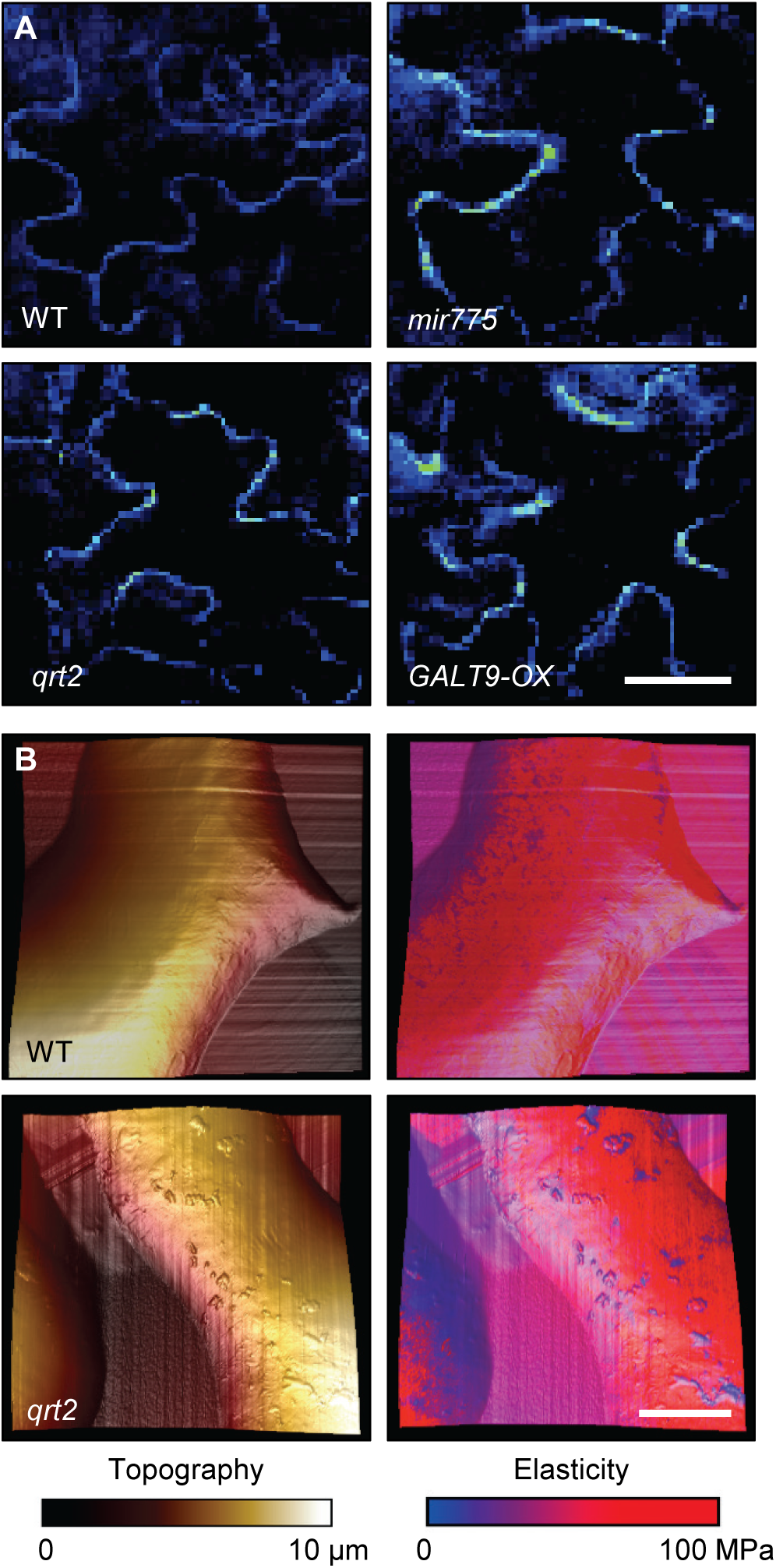
Analysis of the *qrt2* Mutant Defective in Pectin Turnover. (**A**) Examination of cell wall pectin by Raman microscopy. Cotyledon cells of seven-day-old wild type, *mir775*, *GALT9-OX*, and *qrt2* seedlings were imaged for pectin. Bar, 20 μm. (B) Topography of the wild type and *qrt2* cotyledon epidermal cells mapped by AFM (left) and topography overlaid with elasticity (right). Bar, 5 μm. Supports Figures 8 and 9 in the main manuscript.

**Supplemental Figure 11.**
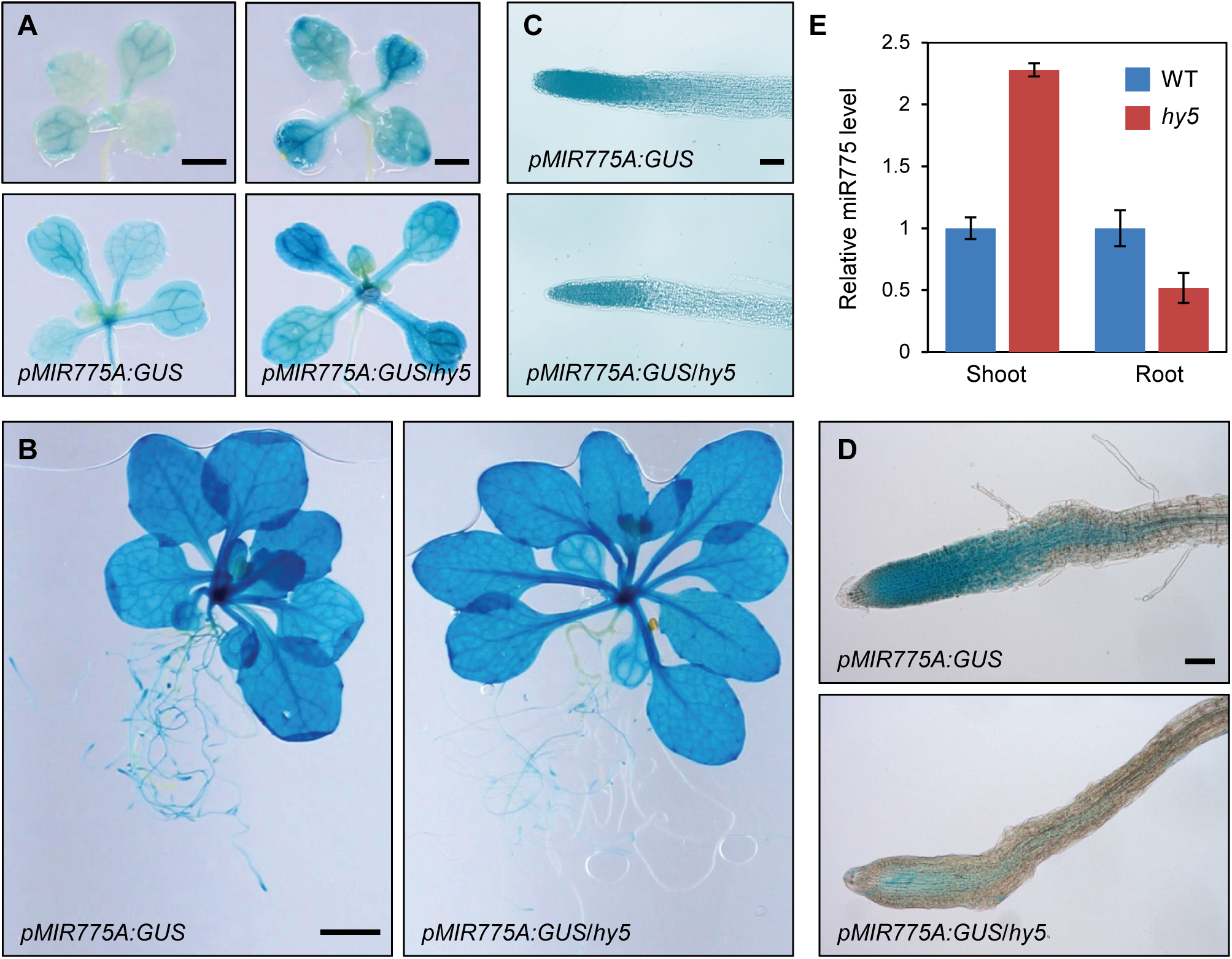
H*Y*5 Differentially Regulates *MIR775A* in the Shoot and the Root. (**A**) GUS staining for *pMIR775A* activities in the wild type and *hy5-215* backgrounds. Ten- (left) and 12-day-old (right) *pMIR775A:GUS* and *pMIR775A:GUS*/*hy5-215* seedlings (right) were stained for GUS activity. Bars, 1 mm. (**B**) The *pMIR775A:GUS* and *pMIR775A:GUS*/*hy5* adult plants with approximately ten true leaves were stained for GUS activity. Bar, 2 cm. (**C-D**) Root tips of *pMIR775A:GUS* and *pMIR775A:GUS*/*hy5* at the seedling (C) and adult (D) stages were compared for GUS activity. Bars, 50 μm. (**E**) Quantitative analysis of relative miR775 levels separately in the shoot and the root of wild type and *hy5-215* seedlings by RT-qPCR. Data are means ± SD from three technical replicates. Supports Figure 10 in the main manuscript.

**Supplemental Figure 12.**
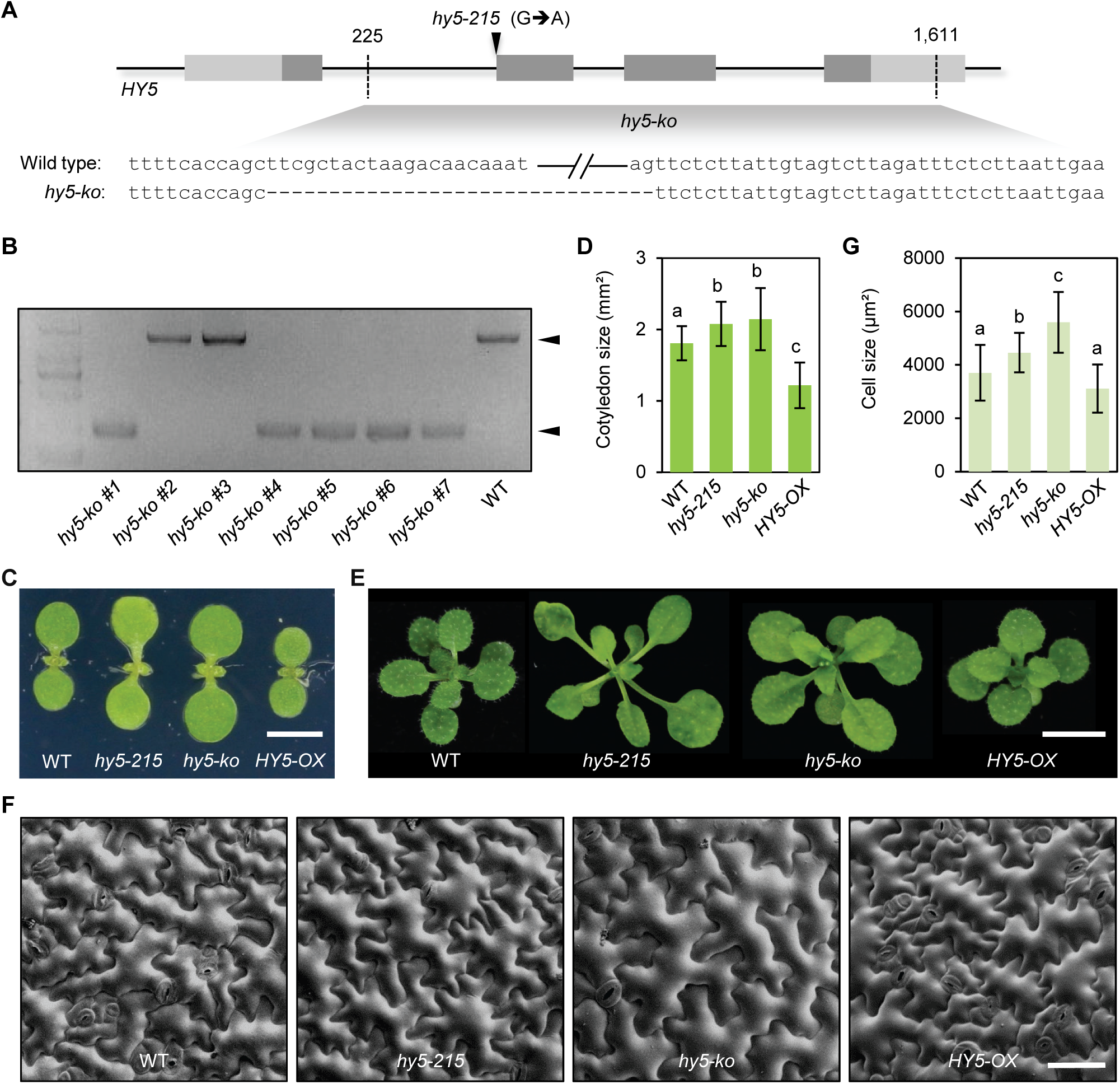
Generation and Characterization of Mutants for *HY5*. (**A**) Scheme for generating the *hy5-ko* allele using CRISPR/Cas9. Two sgRNAs are designed to create paired cleavage sites resulting in a 1,386 bp deletion. The *hy5-215* allele harbors a point mutation near the end of the first intron that interferes splicing. (**B**) Genotyping result with PCR products according to the wild type and deletion alleles indicated. Lines #4 and #5 were selected for subsequent analyses. (**C-D**) Morphological comparison and quantification of cotyledon size. Data are mean ± SD from 10 individual seedlings. Different letters denote genotypes with significant difference (Student’s *t*-test, *p* < 0.05). Bar, 2 mm. (**E**) Morphological comparison of adult plants. Bar, 5 mm. (**F-G**) SEM analysis of the cotyledon epidermal cells. Bar, 100 μm. (**G**) Quantification of the cotyledon epidermal cell size. Data are mean ± SD from 100 individual cells from three seedlings. Different letters denote genotypes with significant difference (Student’s *t*-test, *p* < 0.05). Supports Figures 11 and 12 in the main manuscript.

